# Neurodevelopmental Patterns of Early Postnatal White Matter Maturation Represent Distinct Underlying Microstructure and Histology

**DOI:** 10.1101/2022.02.11.480169

**Authors:** Arash Nazeri, Željka Krsnik, Ivica Kostović, Sung Min Ha, Janja Kopić, Dimitrios Alexopoulos, Sydney Kaplan, Dominique Meyer, Joan L. Luby, Barbara B. Warner, Cynthia E. Rogers, Deanna M. Barch, Joshua S. Shimony, Robert C. McKinstry, Jeffrey J. Neil, Christopher D. Smyser, Aristeidis Sotiras

## Abstract

During the early postnatal period, cerebral white matter undergoes rapid maturation through a complex series of interrelated cellular and histogenetic processes. Accurately quantifying these processes is important for improving understanding of early brain development, developmental abnormalities related to prematurity, and neurodevelopmental diseases. Past efforts have used magnetic resonance imaging (MRI) to track these developmental processes *in vivo*. However, most previous studies have relied on single imaging modality data and have often been limited by small samples and analytics that do not evaluate complex multivariate imaging patterns. Here, we applied an advanced unsupervised multivariate pattern analysis technique, non-negative matrix factorization (NMF), to T_2_w/T_1_w signal ratio maps from a large cohort of newborns (Developing Human Connectome Project [dHCP], n=342), revealing patterns of synchronous white matter maturation. These patterns showed divergent age-related maturational trajectories and differential susceptibility to premature birth, which were replicated in an independent large sample of newborns (Early Life Adversity, Biological Embedding, and Risk for Developmental Precursors of Mental Disorders [eLABE], n=239). Furthermore, we showed that T_2_w/T_1_w signal variations in white matter maturational patterns are explained by differential contributions of white matter microstructure indices (i.e., free water content and neurite density index) derived from neurite orientation dispersion and density imaging (NODDI) modeling of diffusion-weighted MRI. Finally, we demonstrated how white matter maturation patterns relate to distinct histological features by comparing our findings with postmortem late fetal/early postnatal brain tissue staining. Together, these results delineate a novel MRI representation of white matter microstructural and histological reorganization during the early postnatal development.

## Introduction

The late fetal and early postnatal period is a critical stage of human brain development, during which the cerebral white matter undergoes complex changes (Barkovich and Barkovich, 2019; Gilmore et al., 2018; Vasung et al., 2019), characterized by myelination of the major white matter bundles, maturation of the extracellular matrix (ECM), and continued dissolution of the transient laminar organization of the fetal period, such as disappearance of the subplate zone at the white matter-cortex interface (Barkovich and Barkovich, 2019; Kinney and Volpe, 2018; Kostovic et al., 2014). However, early postnatal white matter maturation may be disrupted by preterm birth interrupting normal fetal brain development (Mathur et al., 2010; Volpe, 2009) or altered by other environmental or genetic factors, leading to neurological and psychiatric disorders later in life (Guma et al., 2019; Hazlett et al., 2017; Marín, 2016). Thus, understanding patterns of typical white matter development during this period is necessary for accounts of both healthy brain maturation and neurodevelopmental disorders.

A growing body of studies have investigated white matter maturation using high quality magnetic resonance imaging of newborns. Volumetric studies have demonstrated a steep white matter volume expansion in the early postnatal period, and a relatively decreased white matter volume in preterm infants (Alexander et al., 2019; Makropoulos et al., 2016). Diffusion tensor imaging (DTI) studies have further revealed substantive changes in white matter microstructure including decrease in white matter diffusivity and increase in fractional anisotropy during this period (Dean et al., 2017; Dubois et al., 2014; Gilmore et al., 2007; Mukherjee et al., 2001, 2002), along with altered diffusion metrics in preterm infants (Brenner et al., 2021; Knight et al., 2018). Additionally, the evolution of MRI signal on T_2_-weighted (T_2_w) and T_1_-weighted (T_1_w) images have been used to track early postnatal white matter microstructural changes, such as myelination and ECM maturation (Corbett-Detig et al., 2011; Kostović et al., 2002; Kostovic et al., 2014; O’Muircheartaigh et al., 2020; Paredes et al., 2016; Pittet et al., 2019; Wang et al., 2019; Widjaja et al., 2010; Zunic Isasegi et al., 2018).

Although Magnetic Resonance Imaging (MRI) studies have improved our understanding of early postnatal white matter development, several limitations are notable within the current literature in relation to both imaging modalities and analysis. Most studies have relied on single modality data despite the importance of cross-examining findings with other neuroimaging modalities and brain histology to better shed light on underlying mechanisms. While white matter maturational processes tend to occur in radially-organized periventricular, intermediate, and superficial white matter compartments (Kostović et al., 2014; Kostovic et al., 2014; Pittet et al., 2019), many studies have focused on white matter tracts (Dean et al., 2017; Gilmore et al., 2007). Therefore, tract-based approaches may not be able to fully capture the compartmental maturation of the white matter. Additionally, region-of-interest approaches (Kunz et al., 2014) are limited to the *a priori* selection of regions, which restricts their ability in identifying and characterizing diffuse developmental processes that take place across multiple brain areas. Finally, past accounts have frequently been limited by small sample sizes and analytics that do not evaluate complex multivariate imaging patterns.

To overcome previous limitations, we capitalized on a large sample of newborns, the Developing Human Connectome Project (dHCP) (Makropoulos et al., 2018). Toward better characterizing critical postnatal white matter developmental processes, which result in opposing effects on T_1_w and T_2_w signal intensity (decreased intensity on T_2_w and increased intensity on T_1_w images) (Corbett-Detig et al., 2011; Kostović et al., 2002; Kostovic et al., 2014; O’Muircheartaigh et al., 2020; Paredes et al., 2016; Pittet et al., 2019; Wang et al., 2019; Widjaja et al., 2010; Zunic Isasegi et al., 2018), and enhancing contrast-to-noise ratio (Glasser and Van Essen, 2011; Lee et al., 2015), we calculated T_2_w / T_1_w signal intensity ratio maps. We further leveraged advances in unsupervised multivariate pattern analysis to analyze white matter T_2_w/T_1_w ratio maps. Specifically, we delineated spatial patterns of white matter maturation using non-negative matrix factorization (NMF), which provides a parts-based representation of complex high dimensional datasets (Brunet et al., 2004; Lee and Seung, 1999; Wu et al., 2016) and has recently been adapted for neuroimaging data (Sotiras et al., 2015, 2017).

We hypothesized that early postnatal white matter changes occur in concert with distinct spatial patterns that follow laminar compartmental white matter reorganization. We further predicted that these spatial patterns would show distinct age trajectories of MRI signal evolution, exhibit different susceptibilities to preterm birth, and represent distinct underlying white matter microstructure and histology. As hypothesized, the NMF-derived set of coordinated white matter maturation patterns exhibited a radially-distributed compartmental organization with respect to the lateral ventricles and the cortical ribbon, and showed distinct maturational trajectories. We replicated the differential age-related rate of decline in T_2_w/T_1_w signal in the identified white matter patterns in an independent large imaging cohort of newborns, part of the Early Life Adversity, Biological Embedding, and Risk for Developmental Precursors of Mental Disorders (eLABE) study. Moreover, given the complexity and regional variations of white matter signal changes, we further hypothesized that distinct tissue microstructural properties contribute to white matter development. To this end, we used neurite orientation dispersion and density imaging (NODDI) modeling of diffusion-weighted MRI (from the dHCP study) to determine the microstructural underpinnings of white matter maturational patterns (Zhang et al., 2012a). We observed that changes in white matter free water content and neurite density differentially contribute to patterns of T_2_w/T_1_w signal changes. Finally, we compared our findings with postmortem late fetal/early postnatal brain specimens from the Zagreb Collection of Human Brains (Hrabač et al., 2018) and showed how white matter maturation patterns relate to distinct histological features.

## Results

### Description of dHCP and eLABE participants

After quality control of neuroimaging data, T_2_w and T_1_w images from 342 newborns from the dHCP cohort were included in the main study (age at scan: 35-45 weeks postmenstrual, Table 1). Of the 72 preterm-born infants in the dHCP data (born before 37 weeks gestation), 39 underwent MRI at term-equivalent age or older (i.e., at the time that they reached 37 weeks postmenstrual age or later). Quality-controlled MRI data from the eLABE cohort (n=239; age at scan: 38-44 weeks postmenstrual, Table 1) were used for replication, which included data from 38 preterm-born infants. Of note, all participants in the eLABE study were imaged at term-equivalent age or older.

**Table 1.**
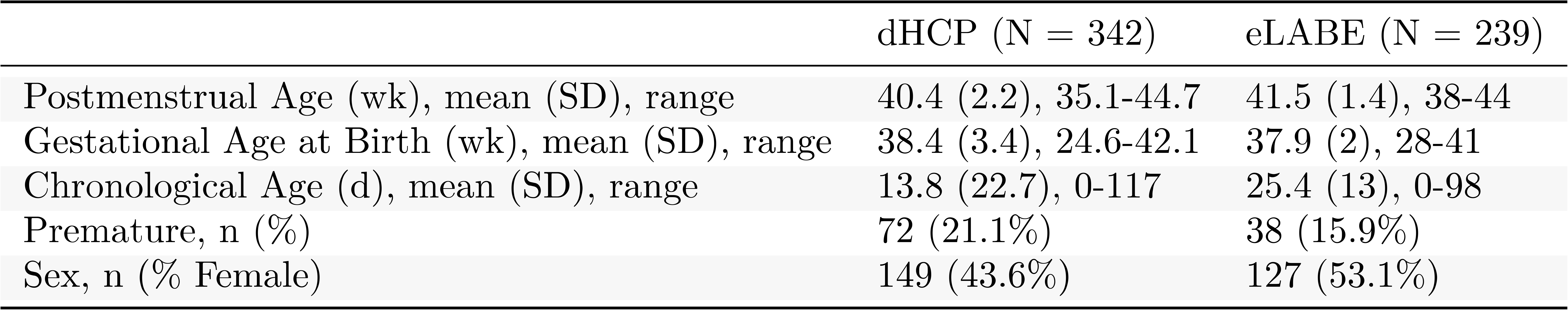
Demographic characteristics of the newborns in the dHCP and eLABE cohorts (the subsets that were used in the study).

### Postmenstrual age and prematurity strongly impact white matter T_2_w/T_1_w signal ratio

We first sought to investigate whether the white matter T_2_w/T_1_w ratio is a sensitive marker for early postnatal cerebral white matter development. Accordingly, we examined the association of mean white matter T_2_w/T_1_w with postmenstrual age and prematurity using generalized additive models (GAMs). Results revealed that mean white matter T_2_w/T_1_w was significantly associated with age and preterm birth in the dHCP cohort. Specifically, mean white matter T_2_w/T_1_w values nonlinearly decreased with age (*R^2^*= 0.30; effective degrees of freedom [edf]= 2.85; *p* <2×10^-16^; Figure 1). There was no significant sex effect or sex-by-age interaction effect on mean white matter T_2_w/T_1_w ratio. Among the term-equivalent or older infants (n_Term_= 270, n_Preterm_= 39), preterm birth was associated with a higher mean white matter T_2_w/T_1_w signal ratio (adjusted for postmenstrual age: t= −8.5, *p*= 5.4×10^-16^; Figure 1). Next, we repeated these analyses in the eLABE cohort, obtaining consistent results. We observed a significant linear age-related decline in mean white matter T_2_w/T_1_w signal ratio (*R^2^*= 0.22, edf= 1, *p*= 9.3×10^-15^) and a trend towards a higher T_2_w/T_1_w signal ratio in preterm newborns (adjusted for postmenstrual age: t= −1.8, *p*= 0.06; n_Term_= 201, n_Preterm_= 38). The absence of images from younger newborns (<38 weeks postmenstrual) in the eLABE study has likely affected the analysis power to identify nonlinear age effects in this cohort. Moreover, the weaker effect observed for prematurity in the eLABE study likely stems from the older gestational age at birth among its preterm newborns (mean [standard deviation, SD] postmenstrual age of preterm newborns in eLABE= 34.3 [2.2] weeks vs. dHCP= 31.5 [3.6] weeks; *p*= 8.2×10^-5^).

**Figure 1.**
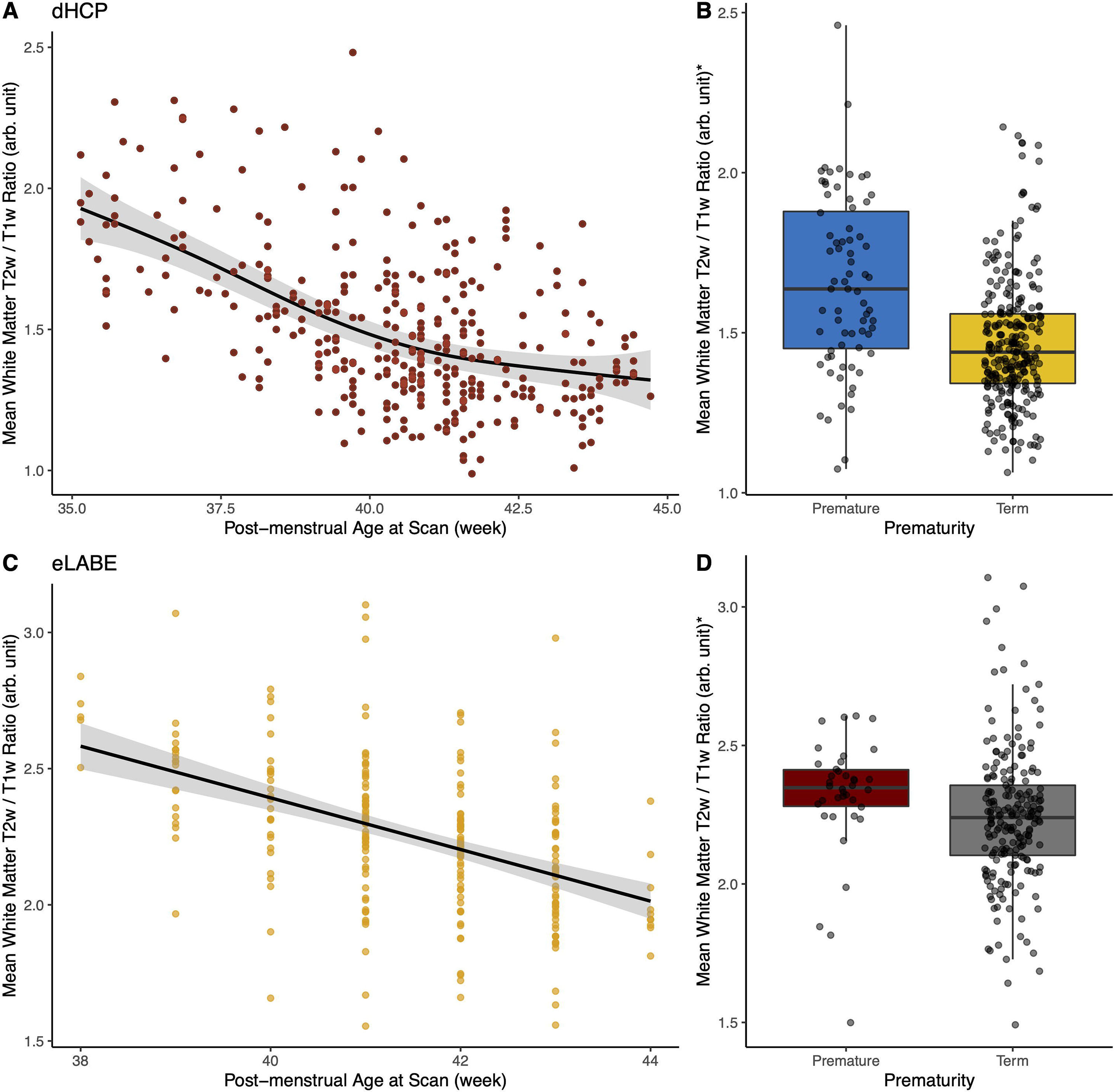
Effects of age and preterm birth on mean white matter T_2_w/T_1_w signal intensity ratio. (A) Nonlinear decrease in white matter T_2_w/T_1_w signal intensity ratio with age in the dHCP cohort. (B) Effect of preterm birth on white matter T_2_w/T_1_w signal intensity ratio in the dHCP cohort. (C) Decrease in mean white matter T_2_w/T_1_w signal intensity ratio with age in the eLABE cohort. A linear model was fitted given that the majority of the subjects in the eLABE cohort fell into a narrower range of age (39-43wk postmenstrual age). (D) Effect of preterm birth on white matter T_2_w/T_1_w signal intensity ratio in the eLABE cohort. **\***Age-adjusted mean white matter T_2_w/T_1_w signal intensity ratio values.

### NMF identifies reproducible and hierarchical early postnatal white matter developmental patterns

We next sought to investigate whether white matter matures through the development of spatially heterogeneous, yet regionally coordinated patterns (Figure 2). To this end, we applied NMF to the white matter T_2_w/T_1_w ratio maps from the dHCP cohort. To identify the most reproducible and reliable solution with the least reconstruction error, we examined multiple NMF solutions ranging from 2 to 20 patterns. Split-half reproducibility analysis, as quantified using the Adjusted Rand Index (ARI)(Hubert and Arabie, 1985), demonstrated that solutions were highly stable, emphasizing the robustness of the identified patterns. However, reproducibility and reliability were not uniform. Clear peaks of reproducibility and reliability occurred for the 3-pattern (mean ARI: 88.2%, SD=0.5%) and 9-pattern (mean ARI: 91.1%, SD=0.7%; Figure 2B) solutions, which were explored further.

**Figure 2.**
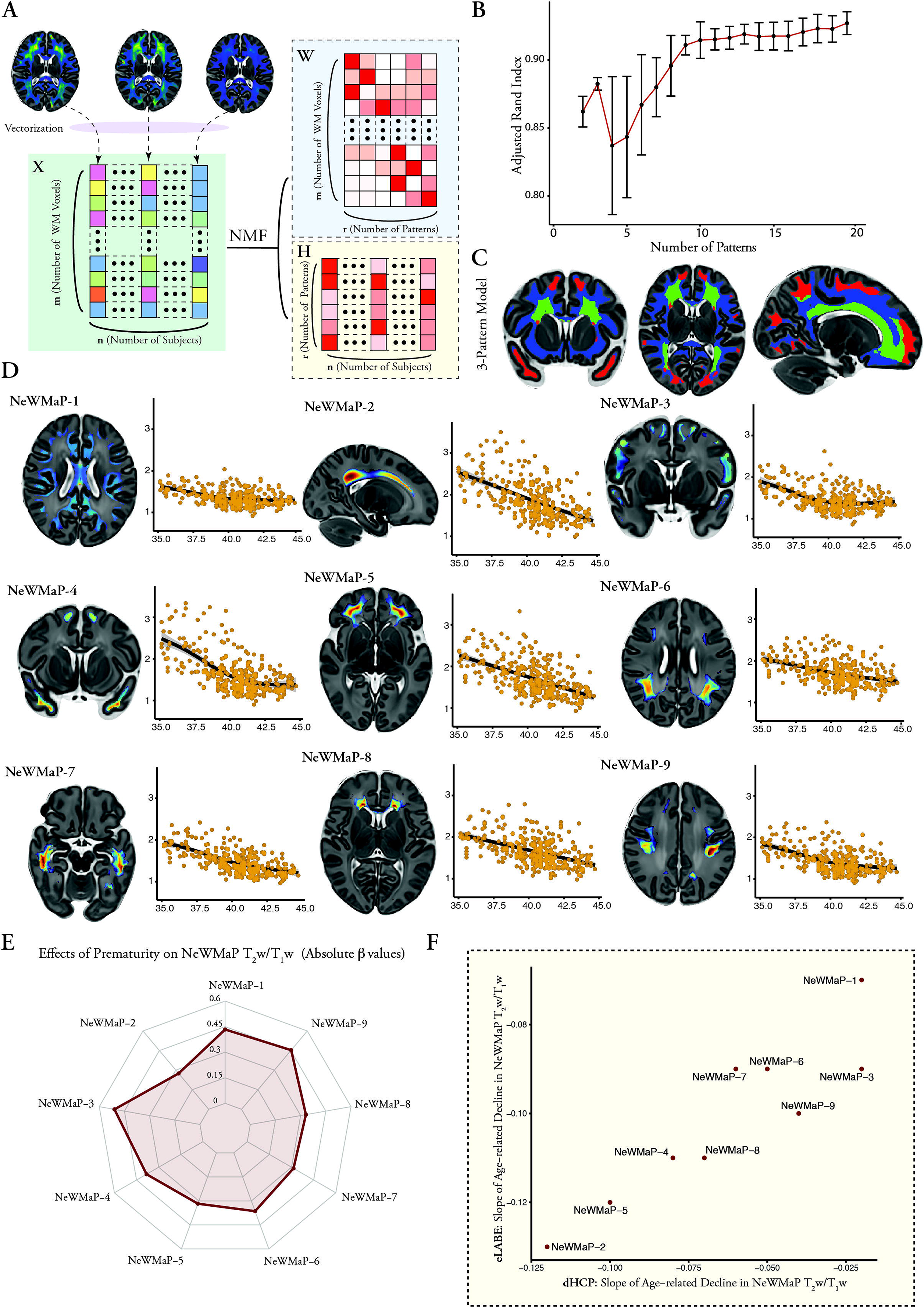
Non-negative matrix factorization (NMF) of T_2_w/T_1_w ratio maps. (A) An overview of T_2_w/T_1_w ratio image data NMF. Three-dimensional white matter T_2_w/T_1_w ratio maps are initially vectorized and then concatenated into matrix **X** (each column representing a subject; each row representing a voxel in the template space). NMF then decomposes the matrix **X** into: (i) matrix **W**: the factor representations (each column representing a factor; each row representing a white matter voxel in the template space); (ii) matrix **H**: the subject-specific loading coefficients for each factor (each column representing a subject; each row representing a factor). (B) Split-half reproducibility analysis with bootstrapping of NMF solutions at multiple resolutions ranging from 2 to 20 white matter factors (mean and standard deviation of adjusted rand index over 20 bootstrap resamples). (C) Maps of the NeWMaPs from the three-factor NMF model. (D) Maps of the NeWMaPs from the nine-factor NMF model and corresponding age-related changes in mean T_2_w/T_1_w signal ratio in these NeWMaPs. NeWMaP-2, NeWMaP-5, and NeWMaP-4 showed the highest early postnatal T_2_w/T_1_w values, while NeWMaP-1 and NeWMaP-9 showed the lowest early postnatal T_2_w/T_1_w values. (E) Effects of prematurity on NeWMaP T_2_w/T_1_w signal ratio in term-equivalent or older newborns. (F) Age-related decline rates of NeWMaP T_2_w/T_1_w signal ratio in the eLABE study and a comparable subset of the dHCP newborns (linear models; postmenstrual age ≥ 38 weeks).

The 3-pattern solution distinguished patterns by largely following distance from the lateral ventricles. The first pattern comprised periventricular white matter, which further extended into the deep white matter of the frontal and parietal lobes. The second pattern consisted of frontotemporal polar superficial white matter, while the remaining superficial and deep white matter areas formed the third pattern (Figure 2C).

These three patterns were further divided hierarchically as part of the 9-pattern NMF solution (Figure 2D; Figure S2; Table S2). Given that the 9-pattern solution demonstrated a lower reconstruction error than the 3-pattern model (Figure S1), the nine <u>Ne</u>wborn <u>W</u>hite matter <u>Ma</u>turation <u>P</u>atterns (NeWMaPs) were used for downstream analyses. Despite being derived in a completely unsupervised and data-driven manner, NeWMaPs were remarkably symmetric and demonstrated a characteristic distance distribution with respect to the cortex and/or the lateral ventricles (Figure S3, Table S2). Three NeWMaPs localized mainly to the superficial white matter: (i) NeWMaP-1: predominantly sulcal aspects of the superficial white matter throughout the brain in addition to major white matter bundles (e.g., corpus callosum, cingula, and external capsules); (ii) NeWMaP-4: superficial white matter in the temporal poles, frontal poles/superior frontal gyri, precunei, and occipital poles; (iii) NeWMaP-3: all other gyral superficial white matter areas. Three other NeWMaPs corresponded to the periventricular white matter: (i) NeWMaP-2: dorsal frontoparietal periventricular region and centrum semiovale; (ii) NeWMaP-7: occipitotemporal periventricular region; (iii) NeWMaP-8: corpus callosum genu/forceps minor and frontal periventricular region. The last three NeWMaPs were intermediate in distance to the cortex and the lateral ventricles: (i) NeWMaP-5: rostral/orbital frontal intermediate region; (ii) NeWMaP-6: posterior parietal superior longitudinal fasciculus, and parietal and dorsal frontal intermediate region; (iii) NeWMaP-9: superior longitudinal fasciculus, perirolandic white matter surrounding the central sulci and occipital poles.

### NeWMaPs display differential developmental effects

The data-driven NMF approach identified patterns of covariance in this large developmental sample, but did not include information regarding participant postmenstrual age or prematurity. Accordingly, we next investigated the developmental patterns across NeWMaPs using GAMs. While all NeWMaPs demonstrated a significant age-related decline in mean T_2_w/T_1_w signal ratio, their trajectories displayed substantial heterogeneity. The steepest decline was observed in periventricular NeWMaP-2 and the smallest rate of decline was in superficial NeWMaP-1 (Figure 2D, Table S3). Among term-equivalent newborns, preterm birth was associated with a higher mean T_2_w/T_1_w signal ratio in all NeWMaPs, with the strongest effect on superficial NeWMaP-3, followed by intermediate NewMaP-9 and superficial NeWMaP-1 (Figures 2E and S4). Moreover, all three periventricular and two of the intermediate NeWMaPs showed less steep rate of decline in T_2_w/T_1_w signal ratio in premature newborns (age × prematurity interaction effects with nominal *p* <0.05: NeWMaP-2: *p =* 0.028; NeWMaP-5: *p =* 0.014; NeWMaP-6: *p =* 0.028; NeWMaP-7: *p =* 0.025; NeWMaP-8: *p =* 0.0021).

To determine whether similar age trajectories would be found in an independent study, we used the NeWMaPs derived from the dHCP cohort as regions of interest to extract mean T_2_w/T_1_w signal ratios in the eLABE study. Similar to dHCP results, all NeWMaPs demonstrated a significant age-related decline in T_2_w/T_1_w signal ratio (Figure S5). To directly compare dHCP and eLABE results, we calculated the correlation between the rates of age-related decline in T_2_w/T_1_w signal ratio across all NeWMaPs. To eliminate the effect of postmenstrual age differences between the two studies, the correlation was calculated between the eLABE sample (all postmenstrual ages are ≥38 weeks) and a subset of dHCP newborns (n=296), which was obtained by excluding participants with postmenstrual age <38 weeks. We found a high correlation between the rates of age-related decline in T_2_w/T_1_w signal ratio of the NeWMaPs in the dHCP and eLABE studies (Figure 2F, Spearman ρ: 0.92, *p* = 0.001; Table S4).

### Free water content and neurite density differentially contribute to NeWMaP T_2_w/T_1_w signal ratio

Given the heterogeneity in T_2_w/T_1_w signal ratio across the NeWMaPs, we hypothesized that distinct white matter tissue microstructural properties contribute to the observed signal changes during early postnatal white matter maturation. Hence, we investigated whether neurodevelopmental patterns derived from T_2_w/T_1_w signal ratio are linked to differences in tissue microstructure indices derived independently from diffusion-weighted MRI. To this end, we fitted the NODDI model to the available multi-shell diffusion MR images in the dHCP dataset(Zhang et al., 2012a) (n=323, Table S1). Unlike DTI, the NODDI model can disentangle compartmental diffusion properties in complex tissue microenvironments, such as early postnatal white matter(Zhang et al., 2012b). Capitalizing on this property, we estimated tissue compartments and generated free water, neurite density, and orientation dispersion maps. We additionally generated weekly average NODDI parameter maps in the template space. This allowed us to visually compare the spatiotemporal patterns of microstructural variations with respect to NeWMaPs. The NeWMaPs were well-demarcated on free water and/or NDI maps (Figure 3A). Furthermore, the application of NeWMaP masks to the weekly average NODDI parameter maps revealed that NeWMaPs largely corresponded to distinct distributions of white matter free water and neurite density (Figure S6).

**Figure 3.**
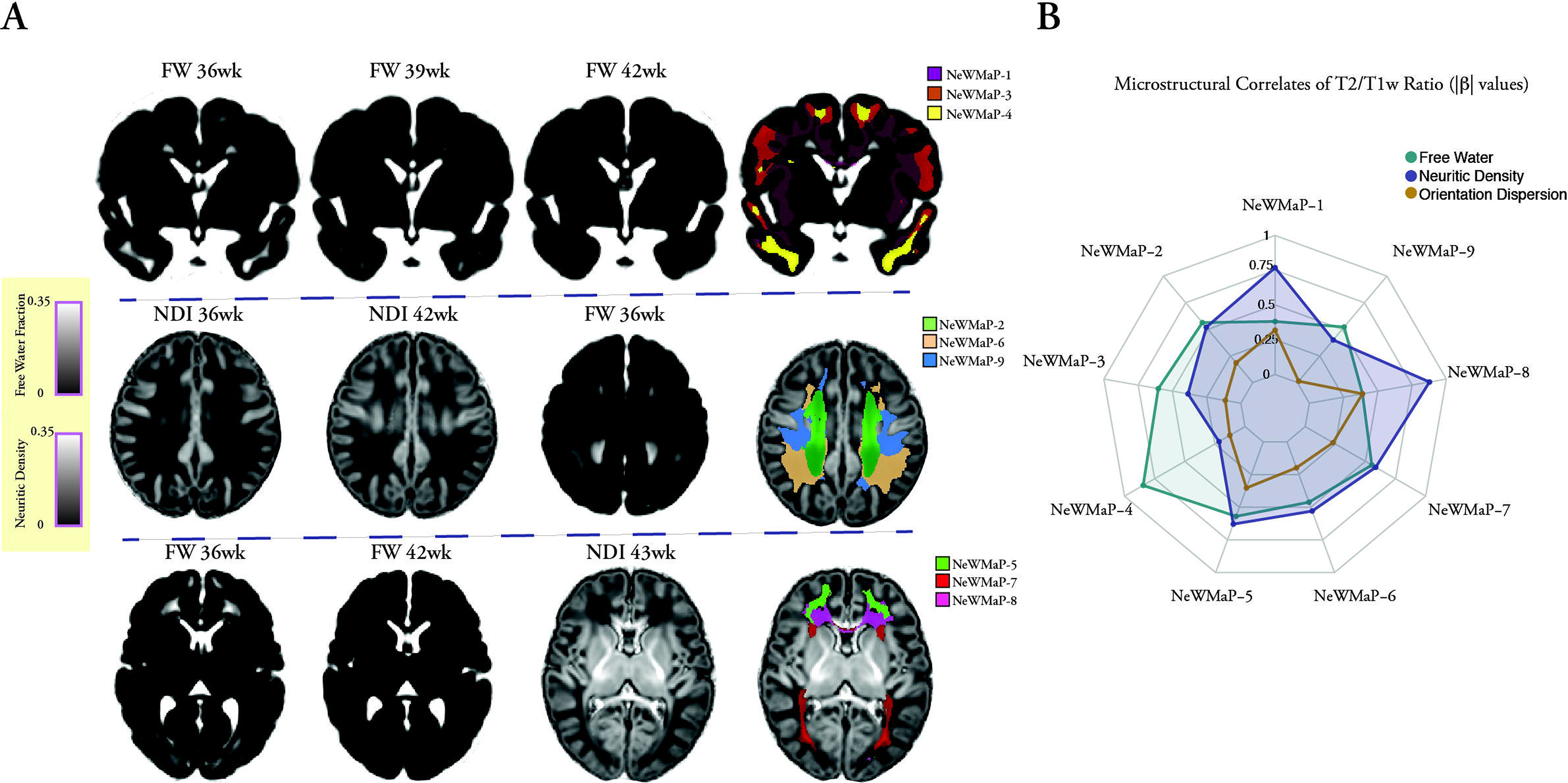
Microstructural correlates of the NeWMaPs. (A) Free water, neurite density, and orientation dispersion index (not shown) maps were generated by fitting the NODDI model into the multi-shell diffusion MR images. Weekly average free water and neurite density maps demonstrate patterns of age-related decrease in free water content and increase in neurite density in the early postnatal period. NeWMaPs largely corresponded to distinct tissue microstructure and were well-demarcated on free water and/or neurite density maps. For instance, unlike NeWMaP-1, the other two superficial white matter NeWMaP-3 and NeWMaP-4 showed relatively high free water contents (NeWMaP-4 demonstrating the highest values). Dorsal frontoparietal periventricular NeWMaP-2 was clearly demarcated from the parietal NeWMaP-6 and frontoparietal NeWMaP-9 on free water and neurite density maps. Occipital portion of the periventricular NeWMaP-7 corresponded to the high neurite density optic radiations. (B) Spiderplot showing effects of free water content, neurite density, and orientation dispersion index on T_2_w/T_1_w from each NeWMaP.

To further investigate links between white matter T_2_w/T_1_w signal ratio and diffusion-derived tissue microstructure indices, we applied NeWMaP masks to individual free water, neurite density, and orientation dispersion maps. This allowed us to quantify the average value for each of these indices within each NeWMaP at the individual level. We used the derived measures as explanatory variables in linear models that predict mean T_2_w/T_1_w signal ratio to assess the respective contributions of white matter microstructure indices to the T_2_w/T_1_w signal ratio in each NeWMaP. Higher free water content was associated with a higher T_2_w/T_1_w signal ratio, while higher neurite density was associated with a lower T_2_w/T_1_w signal ratio in all NeWMaPs. Higher orientation dispersion was associated with a higher T_2_w/T_1_w signal ratio in some NeWMaPs (mainly among the NeWMaPs with higher neurite density contributions; Table S5). Across all NeWMaPs, NODDI indices together explained the majority of white matter T_2_w/T_1_w signal ratio variance (*R^2^*: 0.50-0.85, Table S5). However, the degree of contribution from each tissue microstructure index was different. While neurite density showed a dominant effect on periventricular NeWMaP-8 and superficial NeWMaP-1 T_2_w/T_1_w signal ratio, free water had a dominant effect on the superficial NeWMaP-4 T_2_w/T_1_w signal ratio (Figure 3; Table S5). Free water content significantly decreased with age, while neurite density increased with age across all NeWMaPs (Figures S7 and S8). Similarly, prematurity was associated with significantly higher free water content (except for NeWMaP-5) and lower neurite density in all NeWMaPs (except for NeWMaP-7 and NeWMaP-8; Figure S9, Table S6). These results together with the heterogeneity of developmental effects suggest that different NeWMaPs may be linked in part to distinct neurodevelopmental and histological processes, which we examined next.

### NeWMaPs correspond to late fetal/early postnatal histological reorganization

The hallmarks of late fetal/early postnatal white matter development are the dissolution of subplate remnant, and the maturation of myelinated fibers and astroglial and axonal arrangements. These processes can be tracked through qualitative histopathological analysis of postmortem brain specimens stained for markers of subplate ECM (Alcian-blue), astrocytes and radial glial cells (glial fibrillary acidic protein [GFAP]), myelination (myelin basic protein [MBP]), and axonal maturation (phosphorylated neurofilament proteins; SMI 312). Thus, to investigate the neurobiological underpinnings of NeWMaPs, we examined their histological correlates in appropriately stained late fetal/early postnatal brain specimens from the Zagreb Collection of Human Brains (n=7; Table S7)(Hrabač et al., 2018).

Our first goal was to investigate whether variations in NeWMaP T_2_w/T_1_w signal and NODDI metrics correlate with the ECM content in the subplate remnant. The subplate zone plays a critical role in the development of thalamocortical and corticocortical connections(Kostović, 2020; Kostovic and Judas, 2010) and its disappearance from the interface of the superficial white matter and the cortical ribbon is one of the most drastic changes in the late fetal/early postnatal period. Postmortem histology and fetal/postnatal MRI show that the dissolution of the subplate starts from the sulcal areas, resulting in a thinner “band-like” appearance in sulci and thicker diffuse appearance in gyral white matter in early postnatal period(Kostovic et al., 2014). The subplate neurons and glia are embedded in a highly abundant and hydrophilic ECM, which is selectively stained with Alcian-blue in fetal/postnatal brain(Kostović et al., 2002; Kostovic et al., 2014). Accordingly, we focused our analysis on the superficial NeWMaPs (NeWMaP 1, 3 and 4) and examined coronal brain sections that included these regions and were stained with Alcian-blue. We observed that Alcian-blue staining in the superficial white matter mirrored the observed higher early postnatal T_2_w/T_1_w signal ratio (Figure 2D) and free water content (Figure 3A) in the superficial NeWMaP-3 and NeWMaP-4. Specifically, the *gyral* superficial white matter (NeWMaP-3 and NeWMaP-4) showed higher early postnatal free water contents and more intense and extensive Alcian-blue staining (particularly in the temporal pole [NeWMaP-4] and gyral crowns) compared to the primarily *sulcal* superficial white matter (NeWMaP-1) (Figure 4A-C, Figure S10A). Moreover, deep white matter structures in NeWMaP-1 (e.g., corpus callosum) showed no or minimal Alcian-blue staining.

**Figure 4.**
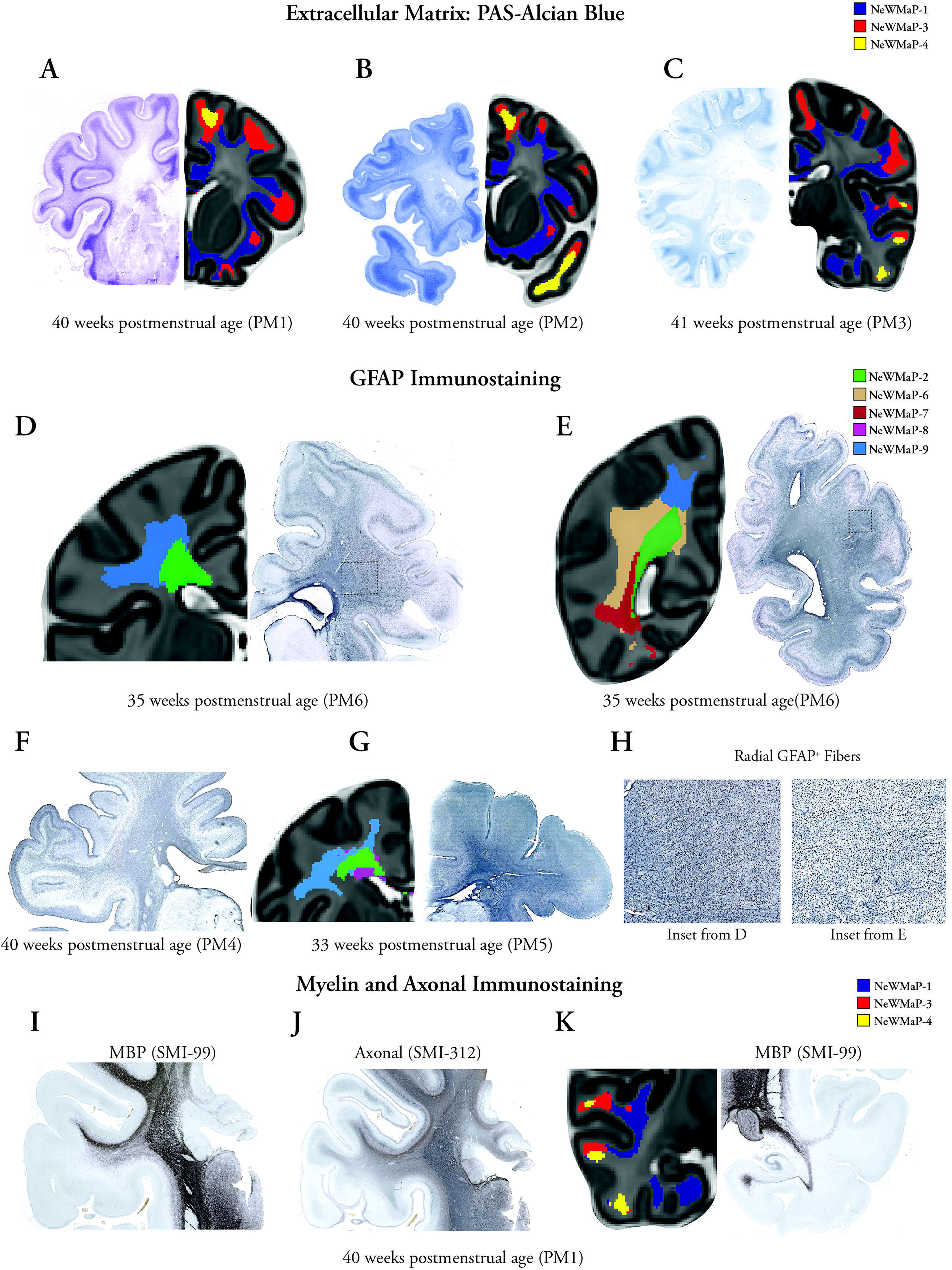
Histological correlates of the NeWMaPs. (A-C) Staining of the extracellular matrix in the subplate remnant with Alcian-blue. Alcian-blue staining of coronal sections through the anterior frontal lobe (A), anterior frontal lobe/temporal pole (B), and frontoparietal junction/posterior temporal lobe (C). Stain vector decomposition was deployed to enhance the Alcian-blue contrast. Corresponding MRI images (T_2_-weighted newborn template) with overlying superficial white matter NeWMaPs (NeWMaP-1, NeWMaP-3, NeWMaP-4) are depicted. Superficial white matter NeWMaP-3 and NeWMaP-4 that localized primarily in the gyri with high early postnatal T_2_w/T_1_w signal ratio and free water content were associated with more intense and thicker Alcian-blue staining. In contrast, sulcal predominant superficial white matter areas corresponding to NeWMaP-1were associated with thinner and less intense Alcian-blue staining. (D-H) Glial fibrillary acidic protein (GFAP) immunohistochemistry of the glial cells. Coronal sections through the frontoparietal junction (D, F, G) and the mid-parietal lobe (E). (D-G) Dorsal frontoparietal periventricular NeWMaP-2 is demarcated from the surrounding intermediate white matter compartment NeWMaP-6 and NeWMaP-9 by its intense GFAP immunoreactivity. (D) Stain decomposition resolves NeWMaP-2 from the surrounding white matter. (H) A distinct feature of the intermediate white matter compartment areas corresponding to NeWMaP-6 and NeWMaP-9 was the presence of radially oriented GFAP-reactive fibers, which likely represent remnants of the radial glial cells. (I-K) Different patterns of myelination and axonal maturation are seen across the NeWMaPs. (I, J) Unlike intermediate white matter compartment NeWMaP-9, periventricular NeWMaP-2 shows no evidence of advanced myelination (no MBP immunoreactivity) and less axonal maturation (less SMI 312 immunoreactivity). (K) Medial temporal lobe (parahippocampal gyrus and entorhinal cortex) and superior temporal gyrus superficial white matter areas corresponding to the NeWMaP-1 showed MBP immunoreactivity, other temporal superficial white matter areas showed no or minimal immunoreactivity. Deep temporal lobe white matter areas showing MBP immunoreactivity either correspond to NeWMaP-7 (not shown) or NeWMaP-1.

Our second goal was to investigate whether differences in cellular elements assessed using GFAP, MBP and neurofilament (SMI 312) staining contribute to the delineation of boundaries between adjacent NeWMaPs. The two main cell types expressing GFAP during human brain development are astrocytes and radial glial cells(Holst et al., 2019). Astrocytes produce ECM components and play various supportive roles crucial for brain homeostasis(Holst et al., 2019; Wiese et al., 2012). Radial glial cells traverse the white matter mantle and contribute to neuronal migration and gliogenesis(Holst et al., 2019; Rash et al., 2019). Examining brain sections immunostained for GFAP, we observed intense GFAP immunoreactivity in the frontoparietal periventricular region corresponding to NeWMaP-2, which is clearly demarcated from the surrounding white matter by its high early postnatal T_2_w/T_1_w signal ratio and free water content (Figure 4D-G). Adjacent dorsal parietal (NeWMaP-6) and frontoparietal (NeWMaP-9) intermediate white matter compartments as well as the occipitotemporal periventricular white matter (NeWMaP-7) demonstrated less GFAP immunoreactivity (Figure 4D-G). On the other hand, a distinct feature of the intermediate white matter compartment areas corresponding to NeWMaP-6 and NeWMaP-9 was the presence of radially oriented GFAP-reactive fibers, which likely represent remnants of the radial glial cells (Figure 4H).

We also examined MBP and SMI 312 immunostained sections and observed different patterns of myelination and axonal maturation across the NeWMaPs. Specifically, white matter areas corresponding to the NeWMaP-9 showed advanced myelination (MBP immunoreactivity) and axonal maturation (SMI 312 immunoreactivity), while the NeWMAP-2 frontoparietal periventricular compartment showed no evidence of myelination in early postnatal brains (Figure 4I, J; Figure S10B). Interestingly, while the medial temporal lobe (parahippocampal gyrus and entorhinal cortex) and superficial white matter of the superior temporal gyrus demonstrated intense MBP immunoreactivity, other areas in the temporal lobe showed either modest (superior temporal gyrus) or no immunoreactivity for MBP. These observations follow the parcellation pattern of the temporal lobe into NeWMaP-1 (medial temporal lobe and superior temporal/Heschl gyrus superficial white matter) and NeWMaP-7 (deep and periventricular white matter), which showed evidence of advanced myelination, as opposed to NeWMaP-3 and NeWMaP-4 (the rest of temporal lobe superficial white matter) (Figure 4K, S10B).

## Discussion

In this study, we capitalized on large developmental cohorts, tailored processing pipelines, advanced multivariate techniques, and flexible modeling to study early postnatal white matter maturation. We showed that T_2_w/T_1_w ratio is a sensitive and robust marker of early postnatal maturation effects. Using unsupervised multivariate pattern analysis, we demonstrated that early postnatal white matter matures in a spatially heterogeneous fashion by integrating highly coordinated units, which demonstrate differential developmental changes. Importantly, we demonstrated that the derived NeWMaPs are not mere statistical constructs but bear biological significance and differences in patterning are driven in part by distinct spatiotemporal white matter microstructure changes and correspond to distinct white matter histology.

We confirmed that by simply dividing T_2_w images by T_1_w images, it is possible to generate quantitative maps of white matter maturation during the early postnatal period. Additionally, by normalizing intensity values, we showed that these maps are sensitive to the early postnatal maturation effects, robust across different MR imaging settings (similar results across two independent large cohorts with different MR imaging platforms, sequences, and resolutions), and reflect spatial variations in white matter microstructure and histological markers of white matter maturation. Our findings demonstrate that various microstructural (i.e., neurite density and free water content) and histological (i.e., myelination, ECM and glial cell pattern and density) factors contribute to the observed T_2_w/T_1_w signal ratio variations in the early postnatal period. Therefore, early postnatal white matter T_2_w/T_1_w signal ratio variations should be interpreted in their spatial contexts, as these variations tend to represent distinct underlying histology in different white matter compartments (e.g., periventricular vs. superficial white matter).

Accordingly, we investigated the spatial variations of T_2_w/T_1_w signal ratio using NMF. We were able to identify reproducible and coherent patterns of white matter signal evolution in the neonatal brain at multiple scales using a data-driven approach. Our approach does not implement any hierarchical constraints, is blind to location in space and is not biased by a preconceived model of white matter development or white matter tracts. However, estimated NeWMaPs at higher resolutions demonstrated increasing differentiation, while largely respecting the boundaries of coarser subdivisions and being predominantly symmetric bilaterally. Importantly, NeWMaPs exhibited a distinctive range of distances from the lateral ventricles and/or the cortex and were not confined to distinct white matter tracts. In addition, the majority of the NeWMaPs were not limited to a specific brain lobe or overlying cortical structure. Instead, NeWMaPs were aligned with white matter compartments(Kostovic et al., 2014), analogous to the laminar and segmental reorganization of the fetal white matter(von Monakow, 1905; Zunic Isasegi et al., 2018). In addition, each NeWMaP demonstrated a characteristic T_2_w/T_1_w signal ratio evolution pattern in early postnatal development. Intriguingly, NeWMaP-1, NeWMaP-3, and NeWMaP-9 that showed the slowest rate of decline in T_2_w/T_1_w signal ratio after 38 weeks postmenstrual age were also the most vulnerable to prematurity, suggesting that these NeWMaPs are at a more developed stage at term and that the late fetal period is more crucial for their maturation. These findings were consistent with earlier reports showing association of preterm birth with increased diffusivity or T_2_ prolongation in earlier-myelinating white matter such as corpus callosum, cingulum, and external capsule (part of NeWMaP-1), corticospinal tract and superior longitudinal fasciculus (part of NeWMaP-9)(Eikenes et al., 2011; Knight et al., 2018).

Importantly, we demonstrated that the derived NeWMaPs reflect distinct underlying microstructural and histological features. We found that low T_2_w/T_1_w signal ratios and high neurite density in white matter structures corresponded to areas of early postnatal myelination(Brody et al., 1987) (e.g., superior temporal gyrus and auditory radiation [NeWMaP-1] and optical radiation [NeWMaP-8]). However, human white matter myelination is a slow and protracted process with its most rapid and dramatic period occurring within the first two years of postnatal life(Kinney and Volpe, 2018; Miller et al., 2012). Therefore, our findings from the early postnatal period capture only a small segment of the extended myelination process. Taken together, these may explain why early postnatal patterns of white matter signal evolution transcended white matter tract definitions and were aligned instead with white matter compartments.

This alignment was further supported by comparing our *in vivo* MRI findings with postmortem late fetal/early postnatal brain sections. Our results suggest that the observed early postnatal high free water content and T_2_w/T_1_w signal in the superficial white matter NeWMaP-3 and NeWMaP-4 correspond to abundant ECM in the subplate remnant. The subplate compartment is the largest transient fetal brain compartment that consists of an abundant hydrophilic ECM, which renders it hyperintense on T_2_w images and contributes to the high free water content estimated from diffusion-weighted MRI(Kostović, 2020; Kostovic et al., 2014). Abundance of ECM in the subplate remnant compartment is a marker of growth of short corticocortical pathways, especially U-shape fibers, which are known to develop postnatally (in contrast to the long corticocortical pathway, which predominantly develop earlier during the intrauterine period)(Burkhalter et al., 1993; Huang et al., 2009; Vasung et al., 2017).

Similar to the subplate remnant, the periventricular crossroad areas, where major association, projection, and commissural white matter fibers intersect are also marked by abundant ECM(Judas et al., 2005; Pittet et al., 2019). Accordingly, the periventricular NeWMaP-2 largely corresponded to the dorsal frontoparietal crossroad area, which had the highest early postnatal T_2_w/T_1_w signal ratio and free water content among the periventricular NeWMaPs. NeWMaP-2 was additionally characterized by intense GFAP staining and absent myelin/axonal maturation staining. Similar to NeWMaP-2, the surrounding frontoparietal intermediate NeWMaPs (NeWMaP-6, NeWMaP-9) also showed high GFAP-reactive glial cell density. However, they were not as GFAP intense and showed more advanced myelination. Moreover, a characteristic finding in these intermediate white matter compartments was the presence of radially oriented GFAP-reactive fibers that likely represent remnants of radial glial cells. Together, these findings suggest that distinct patterns and extents of glial cell GFAP reactivity can be associated with noticeable differences in T_2_w/T_1_w signal ratio. These differences likely arise from the role of radial glial cells and astrocytes in the production and maintenance of the ECM during brain development(Wiese et al., 2012). Periventricular astrocytes and radial glial cells play key roles in fetal brain development, such as neurogenesis and neural migration(Hansen et al., 2010; Nadarajah and Parnavelas, 2002). Neural migration in the frontal lobe has been shown to continue for several months after birth(Paredes et al., 2016; Sanai et al., 2011). However, additional studies are needed to investigate whether postnatal neural migration also occurs in the parietal lobes, potentially contributing to these observed signal variations.

Although this study capitalized on a large sample and advanced methods, certain limitations should be noted. First, the T_2_w/T_1_w signal intensity ratio is an indirect measure of tissue relaxation times. While advances in fast and high-resolution acquisition of these basic MR sequences make it possible to resolve subtle signal changes of the newborn brain in the clinical setting(Kozak et al., 2020), one important technical hurdle is that MR acquisition across multiple sites/scanners inherently introduces undesired variations in image intensity and quality. Intensity harmonization techniques(Wrobel et al., 2020) or relaxometry methods(Leppert et al., 2009) can be utilized to diminish unwanted scanner effects and allow quantitative analysis and individual-level inference for clinical applications. Second, we performed our study in MR images from healthy newborns without incidental neuroimaging findings of clinical significance. It remains to be determined whether this approach can also be applied in the clinical setting to identify abnormal patterns of postnatal white matter signal evolution. Lastly, while we primarily focused on the 9-pattern solution as the simplest solution with high split-half reliability, the optimal number of patterns to describe early postnatal T_2_w/T_1_w signal intensity ratio evolution is likely a function of MR imaging acquisition technique (e.g., image resolution, signal/contrast to noise ratio, T_2_w and T_1_w weighting), age range and demographics of the newborns.

These limitations notwithstanding, the identified early postnatal maturational patterns may provide an alternative to existing tract-based atlas definitions for this critical neurodevelopmental period. By providing a concise representation of early postnatal white matter maturation, these patterns can be exploited to examine the influence of the complex high dimensional genetic and environmental factors on brain development. We further provide normative age trajectories of signal ratio in these NeWMaPs, which can be used to monitor white matter signal evolution during early postnatal development. Additionally, our findings suggest that the maturational patterns are rooted in histological variations across the white matter, largely recapitulating late fetal/early postnatal white matter organization. Taken together, the derived spatial patterns and corresponding developmental trajectories provide important context that can inform neurobiological studies of early postnatal development in health and disease. In conclusion, our study provides insights into early postnatal white matter organization and maturation using high-resolution T_2_w and T_1_w MRI sequences coupled with advanced image processing and analysis techniques. We reveal distinct white matter maturation patterns, which follow distinct developmental trajectories and align well with underlying microstructure and histology.

## Methods

### The Developing Human Connectome Project

Infants were prospectively recruited as a part of the dHCP study, which is an observational cross-sectional open-science project (for details on study design, participant recruitment, and imaging methods, please refer to http://www.developingconnectome.org) (Makropoulos et al., 2018). The study was approved by the National Research Ethics Committee and written consent was obtained from all participating families before imaging.

#### T_1_w and T_2_w MRI Acquisition

366 participants had complete T_1_w and T_2_w datasets (dHCP 2.0 release; 2019). Fourteen infants with poor structural image quality data, incidental findings with potential clinical significance and/or impact on image analysis were excluded from the study (n=342; age at scan: 35-44.5 weeks postmenstrual age; Table 1). All MR images were acquired using a 3T Philips Achieva scanner and a dedicated 32-channel neonatal head coil(Hughes et al., 2017). 2D sagittal and axial turbo spin echo (TSE) sequences were used to acquire both T_1_w and T_2_w images (all with two overlapping stacks of slices with in-plane resolution: 0.8 × 0.8; slice-thickness: 1.6 mm resolution). The images were subsequently corrected for motion and reconstructed at 0.5 × 0.5 × 0.5 mm(Makropoulos et al., 2018).

### T_2_w/T_1_w map generation, preprocessing, and registration

For an overview of image preprocessing and registration please refer to the Figure S9. The dHCP Neonatal Structural Pipeline has been described in detail elsewhere(Makropoulos et al., 2018). After brain extraction, T_2_w-brain images were further segmented into different brain tissues (9 labels including: white matter, cortical gray matter, ventricles) using the Draw-EM algorithm (of note, the white matter label does not include the internal capsule)(Makropoulos et al., 2014). Bias field corrected T_2_w and T_1_w images were used for further analysis. To generate T_2_w/T_1_w maps, the T_2_w images were divided by the T_1_w images for each individual using *fslmaths* (part of FSL(Jenkinson et al., 2012)).

#### Neurodevelopmentally-informed nonlinear registration of the white matter

To preserve the laminar organization of the white matter, we devised a tailored non-linear registration strategy that maintains the underlying white matter topology as a function of distance from the cerebral cortex and the ventricles (Figure S9)(Huang et al., 2006). For each voxel in the white matter, distances to the (i) cortical ribbon (*cortical distance map*) and (ii) to the ventricles (*ventricular distance map*) were calculated using the *distancemap* function in FSL v6.0.0(Jenkinson et al., 2012). These maps complement each other and are more informative at lower values (e.g., closer to the ventricle for the ventricular distance map). Therefore, for both cortical distance and ventricular distance maps all values above 6 mm were set to 6 mm. T_2_w images, cortical, and ventricular distance maps were used to create age-specific templates using the “*antsMultivariateTemplateConstruction2.sh*” script (part of Advanced Normalization Tools [ANTs] repository)(Avants et al., 2011; Tustison and Avants, 2013). This process was repeated for each age group (one template for each week from 35-44wk postmenstrual age). A template-of-templates was generated by applying the same process to the age-specific weekly templates. All templates are provided as supplementary files. The linear and nonlinear transformation files were used to warp images from the native space (T_2_w/T_1_w ratio and white matter mask) to the template space.

#### Preparation for Voxel-based Analysis

To account for residual registration errors, a common white matter mask for voxel-wise analyses was defined as the white matter voxels that were present in more than 90% of the subjects. The non-white matter voxels in any given subject’s nonlinearly warped T_2_w/T_1_w map were filled with the average of the surrounding white matter voxels, weighted by their closeness as determined by using a Gaussian kernel (full-width half-maximum [FWHM]= 1 mm). Finally, spatial smoothing was performed within the white matter mask using the *3dBlurInMask* function (FWHM = 1 mm; part of AFNI v19.2(Cox, 2012)).

### Non-negative matrix factorization

We used NMF to identify patterns where white matter T_2_w/T_1_w ratio covaried consistently across participants. NMF takes as input a tall non-negative data matrix ***X*** (constructed by arraying column-wise vectorized aligned white matter T_2_w/T_1_w ratio maps, ***X*** = 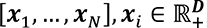, where ***D*** is the number of voxels and ***N*** is the number of samples) and approximates it as a product of two non-negative matrices ***W*** and ***H*** (***X*** ≈ ***WH***, ***W*** ≥ ***0***, ***W*** ≥ ***0***; see Figure 2A for a schematic overview of the NMF procedure). The matrix ***W*** (***W*** = 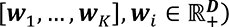 contains in each column one of *K* estimated patterns, where *K* is the user-specified number of patterns. The matrix ***W*** is estimated by positively weighting variables of the data matrix that co-vary in a consistent way across the population and the weight of each variable denotes its relative contribution to the pattern. The matrix ***H*** (***H***^***T***^ = [***h***_1_, …, ***h***_*K*_], ***h***_*i*_ ∈ 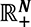) contains subject-specific coefficients for each pattern, which indicate the contribution of each pattern in reconstructing the original T_2_w/T_1_w map. Code for NMF (https://github.com/asotiras/brainparts) adopts orthonormality constraints for the estimated covariation patterns (***W***^***T***^***W*** = ***I***, where ***I*** is the identity matrix) and projective constraints for their respective participant-specific coefficients (***H*** = ***W***^***T***^***X***) (Sotiras et al., 2015; Yang and Oja, 2010). Additional details regarding the implementation of NMF have been presented elsewhere (Sotiras et al., 2015, 2017).

A key difference between NMF and more commonly used methods, such as Principal Component Analysis (PCA) and Independent Component Analysis (ICA), is the non-negativity constraints on the estimated matrices ***W*** and ***H***. Non-negativity constraints have been shown to lead to a parts-based representation (Lee and Seung, 1999), where parts are combined in an additive way to form a whole. This representation yields patterns that enjoy improved statistical power compared with standard mass univariate analyses and are more interpretable and reproducible compared to patterns produced by PCA and ICA (Lee and Seung, 1999; Sotiras et al., 2015).

Consistent with prior studies using this technique (Sotiras et al., 2015, 2017), we ran multiple NMF solutions requesting 2–20 patterns to obtain a range of possible solutions for comparison. We then calculated the reconstruction error for each solution as the Frobenius norm between the data matrix and the NMF approximation and plotted the reconstruction error for all solutions (Figure S1). We additionally performed a split-half reproducibility analysis with bootstrapping with replacement to examine the stability of NMF solutions at multiple resolutions. The optimal number of patterns was chosen based on the solution that best explains the variability in the data (i.e., lower reconstruction error) and reliably shows high out-of-sample reproducibility (reproducibility: mean adjusted rand index [ARI] across 20 bootstraps; reliability: smallest standard deviation of ARI across the bootstraps). ARI provides a measure for set similarity that is adjusted for chance (Hubert and Arabie, 1985). This allows a more balanced comparison between a set of regions when these regions vary in spatial extent. An ARI over 0.75 is deemed excellent (Fleiss et al., 2013).

### Diffusion-weighted MRI and NODDI

Multi-shell diffusion-weighted MR images were collected as a part of dHCP study(Bastiani et al., 2019). Spin-echo echoplanar images were acquired with four different phase encoding directions (Left to Right, Right to Left, Posterior to Anterior, and Anterior to Posterior) and 3 different b-value shells (400, 1000 and 2600 s/mm^2^; Resolution: 1.5×1.5×3 mm^3^, slice overlap: 1.5 mm). A total of 300 volumes were collected with 20 b_0_ images and 64, 88 and 128 unique directions per b-value shell. FSL EDDY was used to correct for motion, motion-induced signal drop-out, and eddy currents(Andersson and Sotiropoulos, 2016). NODDI is a simplified biophysical model that characterizes microstructural tissue contributions to diffusion signal in each voxel into with 3 compartments: intracellular (within neurites), extra-cellular, and free-water compartments(Zhang et al., 2012a). Three main parameters can be derived from this model: 1) free-water isotropic volume fraction; 2) neurite density index, describing intraneurite volume fraction in a given voxel, adjusted for free water (ranging from 0 to 1); 3) the orientation dispersion index (ODI), which characterizes the orientational configuration of neurites in a given voxel (ranging from 0 [highly coherent] to 1 [highly dispersed]). The Microstructure Diffusion Toolbox (MDT v1.2.4; https://github.com/robbert-harms/MDT) was employed for GPU-accelerated fitting of the NODDI model(Harms and Roebroeck, 2018; Harms et al., 2017). NeWMaP masks were transformed into the diffusion space via inverted nonlinear warp fields from the template space to the individual T_2_w space and affine transformation matrix from T_2_w space to diffusion space. We applied the resulting NeWMaP masks in the diffusion space to individual free water, neurite density, and orientation dispersion maps to extract mean values (using the *fslstats* function in FSL).

### The Early Life Adversity, Biological Embedding, and Risk for Developmental Precursors of Mental Disorders (eLABE) study

The eLABE study was approved by the Human Studies Committees at Washington University(Sylvester et al., 2021). Informed consent was obtained from parents of all neonatal participants. Participants were scanned within the first month of life during natural sleep without the use of sedating medications on a Siemens 3T Prisma scanner with a 64-channel head coil. High-resolution 3D T_1_w magnetization-prepared rapid gradient-echo imaging (MPRAGE; TR=2400ms, TE=2.22ms, 0.8mm isotropic) and 3D T_2_w sampling perfection with application-optimized contrasts using different flip-angle evolutions (SPACE; TR=3200 or 4500ms, TE=563ms, 0.8mm isotropic) images were acquired.

T_1_w and T_2_w images underwent correction for gradient nonlinearity-induced and readout distortions(Glasser et al., 2013). The images were then realigned to approximate the MNI152 template orientation (using the *fslreorient2std* script in FSL(Jenkinson et al., 2012)) and denoised (using the *DenoiseImage*(Manjón et al., 2010) function in ANTs). The resulting T_1_w and T_2_w images were coregistered using the *img_reg_4dfp* function (https://4dfp.readthedocs.io/). The Melbourne Children’s Regional Infant Brain atlas Surface (M-CRIB-S) segmentation and surface extraction toolkit was used to generate automated tissue labeling (including white matter, cortical gray matter and ventricles)(Adamson et al., 2020). Finally, the same template creation and nonlinear registration steps that were carried out for the dHCP study were applied to the eLABE images (Figure S9).

### The Zagreb Collection of Human Brains and histological analysis

Late fetal and neonatal post-mortem human brain specimens without macroscopical and microscopical pathological changes were used in the study (n=7; 33-43 postmenstrual age [weeks]; Table S7). Tissue is a part of the Zagreb Collection of Human Brains(Hrabač et al., 2018) obtained during regular autopsies after medically indicated, spontaneous abortions, or after the death of prematurely born or term infants at the clinical hospitals associated to the University of Zagreb, School of Medicine. Tissue sampling was done in agreement with the Declaration of Helsinki (fifth revision, 2000), approved by the IRB of the Ethical Committee at the University of Zagreb, School of Medicine.

Brain tissue was fixed in 4% paraformaldehyde in 0.1M, PBS, pH 7.4, embedded in paraffin, and cut at 20µm thick sections. The coronal sections used in this study were stained by PAS-Alcian blue staining. Immunohistochemistry was performed after deparaffinization of the 20µm thick sections in series of alcohol, 0.3% hydrogen peroxide treatment, and incubation in blocking solution (3% bovine serum albumin, 0.5% Triton x-100, Sigma, St. Louis, MO, USA) in 0.1M PBS, for an hour. Subsequently, sections were incubated with primary antibodies (anti-GFAP, DAKO, z-0334, 1:1000; SMI-99 [anti-myelin basic protein] Biolegend; 808401, 1:1000, anti-SMI-312 [panaxonal anti-neurofilament marker], Biolegend, 1:1000) overnight at room temperature, and following washes, appropriate secondary biotinylated antibodies were applied (Vectastain ABC kit, Vector Laboratories, Burlingame, CA, USA). The sections were developed with 3,3-diaminobenzidine (DAB) with a metal enhancer (Sigma, St. Louis, MO) and coverslipped with Histomount mounting medium (National Diagnostics, Charlotte, NC). All stained slides were imaged using a high-resolution digital slide scanner NanoZoomer 2.0RS (Hamamatsu, Japan). QuPath v0.2.3. was used for visualization of whole-slide images and stain separation with color deconvolution(Bankhead et al., 2017).

### Statistical analysis

All region-of-interest statistical analyses were performed using R v3.6.2(R Core Team, 2018). NMF masks were generated by assigning any given voxel across the white matter to the NMF pattern with the highest loading. NMF masks were registered to the T_2_w/T_1_w ratio maps from the eLABE study and the native diffusion space from the dHCP study. For each NeWMaP, mean T_2_w/T_1_w (both dHCP and eLABE study), free water, neurite density, orientation dispersion values were extracted for each individual using *fslstats* (part of FSL). To model potentially nonlinear effects of postmenstrual age on white matter T_2_w/T_1_w signal ratio, generalized additive models (GAMs) were fit using restricted maximum likelihood (REML, as implemented in *mgcv* package(Wood, 2017)), which produces unbiased estimates of variance and covariance parameters. Generalized linear models were used to assess interaction effects (i.e., sex by age interaction effects) and contribution of NODDI parameters (free water, neurite density, and orientation dispersion index) on T_2_w/T_1_w signal ratio.

## Acknowledgements

AN is a recipient of the Canon Medical Systems USA, Inc./Radiological Society of North America Research & Education (RSNA R&E) Foundation Research Resident Grant (RR1953). AS was partially supported by NIH award R01AG067103. JSS was supported by NIH award P50 HD103525 to the Intellectual and Developmental Disabilities Research Center at Washington University. Data were provided by the developing Human Connectome Project, KCL-Imperial-Oxford Consortium funded by the European Research Council under the European Union Seventh Framework Programme (FP/2007-2013) / ERC Grant Agreement no. [319456]. We are grateful to the families who generously supported this trial. This research was partly funded by the “Research Cooperability” Program of the Croatian Science Foundation funded by the European Union from the European Social Fund under the Operational Programme Efficient Human Resources 2014-2020 PSZ-2019-02-4710 (ZK). Computations were performed using the facilities of the Washington University Center for High Performance Computing, which were partially funded by NIH grants 1S10RR022984-01A1 and 1S10OD018091-01.

