## Supplementary Information for "Neurodevelopmental Patterns of Early Postnatal White Matter Maturation Represent Distinct Underlying Microstructure and Histology"

*Supplementary Information* **-- Neurodevelopmental Patterns of White Matter Transformation during the Early Postnatal Period**

Arash Nazeri, Željka Krsnik, Ivica Kostović, Sung Min Ha, Janja Kopić, Dimitrios Alexopoulos, Sydney Kaplan, Dominique Meyer, Joan L. Luby, Barbara B. Warner, Cynthia E. Rogers, Deanna M. Barch, Joshua S. Shimony, Robert C. McKinstry, Jeffrey J. Neil, Christopher D. Smyser, Aristeidis Sotiras

Table S1. Demographic characteristics of the newborns in the dHCP subsets with term-equivalent or older age-at-scan or with multi-shell data (NODDI).

Table S2. Spatial characteristics of the 9-pattern newborn white matter maturation patterns (NeWMaPs).

Table S3. Age-related changes in NeWMaP T_2_w/T_1_w ratio in the dHCP newborns. The estimated degree of freedom (edf) reflects the degree of non-linearity of a curve in GAM. Age of sharpest decline was calculated for NeWMaPs with edf ≥ 2 (i.e., highly non-linear relationship). PMA: postmenstrual age.

*Table S4.* Age-related decline rates of NeWMaP T_2_w/T_1_w signal ratio in the eLABE study and a comparable subset of the dHCP newborns (linear models; postmenstrual age ≥ 38 weeks).

*Table S5.* Effects (β values) of white matter microstructure indices derived from NODDI on T2w/T1w signal ratio of the NeWMaPs.

* *p* <0.05, ** Significant after correcting for multiple comparisons (*p* <0.0056 for 9 NeWMaPs)

*Table S6.* Effects (β values and standard errors) of prematurity on mean T2w/T1w signal ratio, free water density, and neurite density of the NeWMaPs.

* *p* <0.05, ** Significant after correcting for multiple comparisons (*p* <0.0056 for 9 NeWMaPs)

*Table S7.* Demographic characteristics of subjects from the Zagreb Collection of Human Brains used in the study. **Exact age of this term born subject is unavailable.*

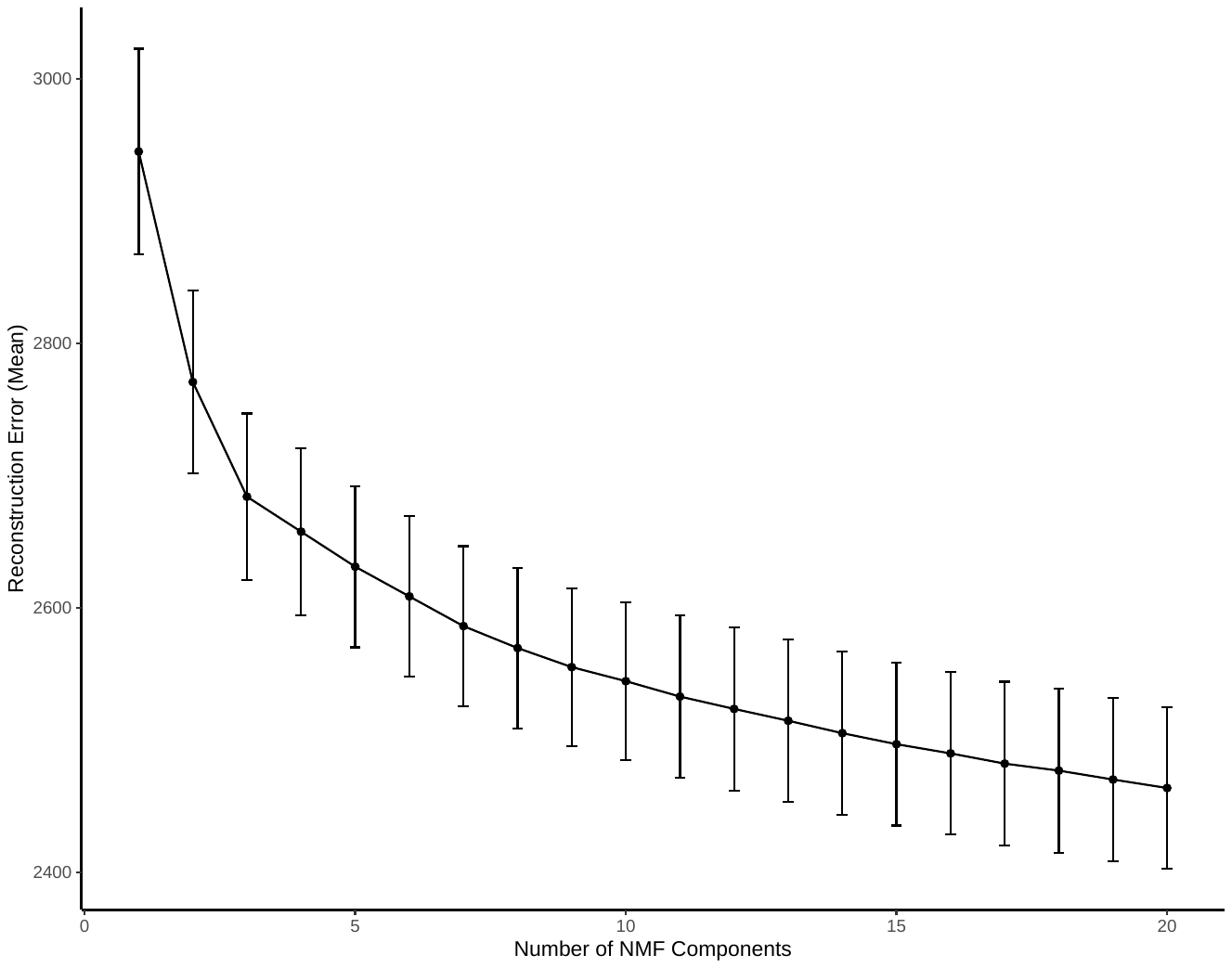

*Figure S1*. Reconstruction errors from split-half bootstrapping of NMF solutions at multiple resolutions ranging from 2 to 20 white matter patterns (mean and standard deviation over 20 bootstrap resamples).

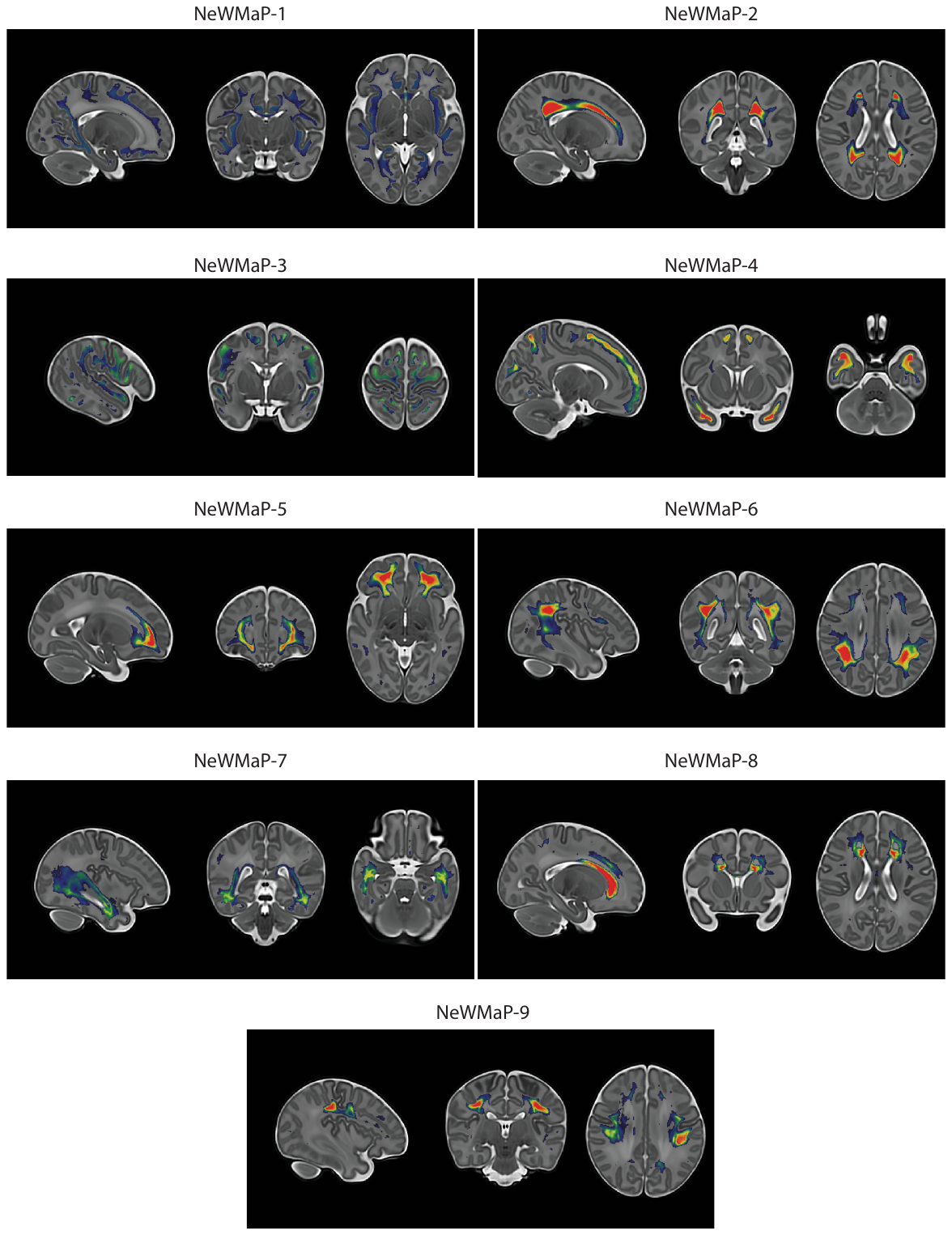
*Figure S2*. Maps of the NeWMaPs from the 9-pattern NMF model in sagittal, coronal, and axial planes.

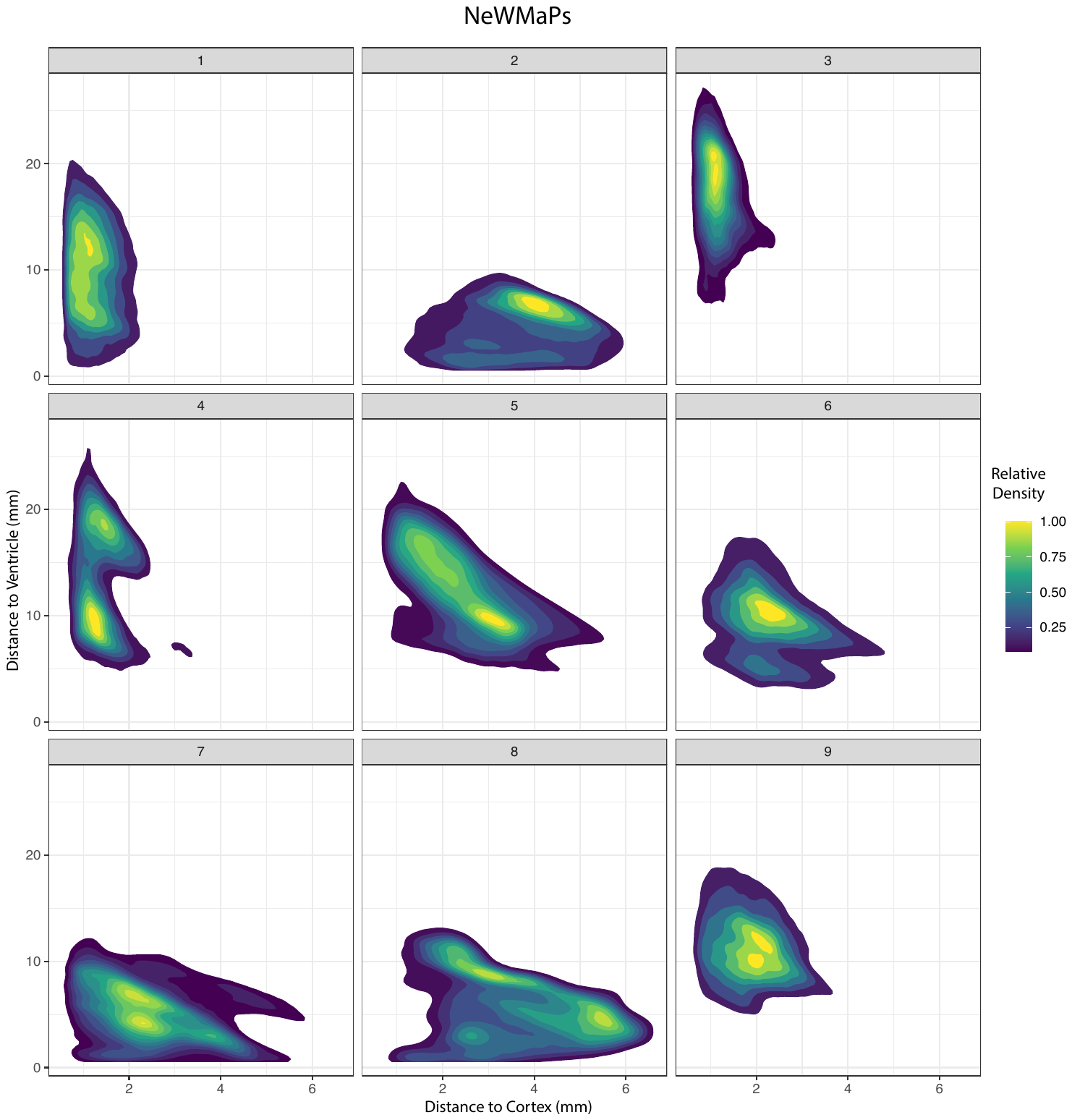
 Figure S3. Two-dimensional density plots demonstrating characteristic distance distributions with respect to the cortex (X-axis) and the lateral ventricles (Y-axis) for each NeWMaP.

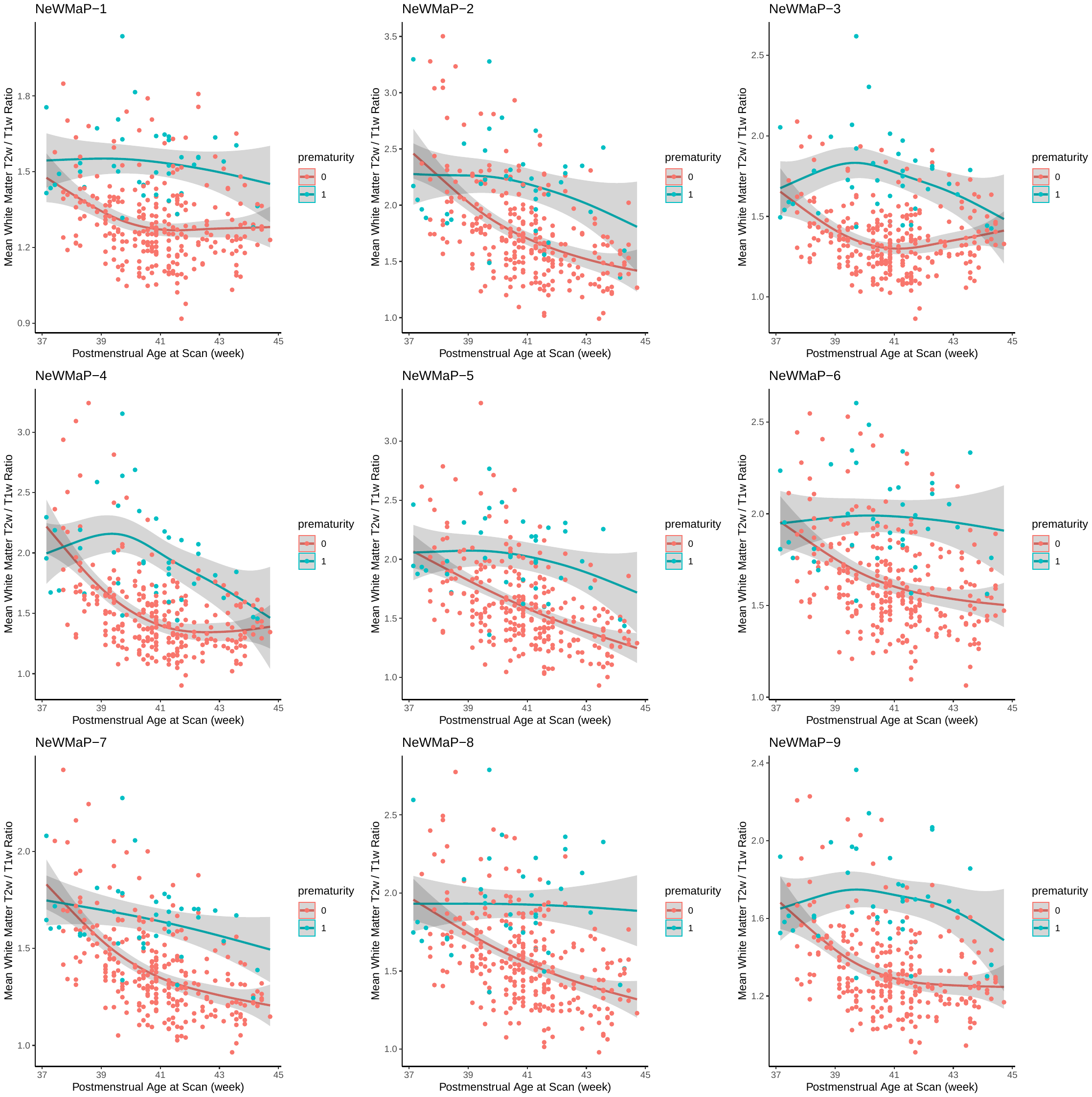

Figure S4. Effect of prematurity on age-related changes in mean NeWMaP T_2_w/T_1_w signal ratio in the dHCP cohort.

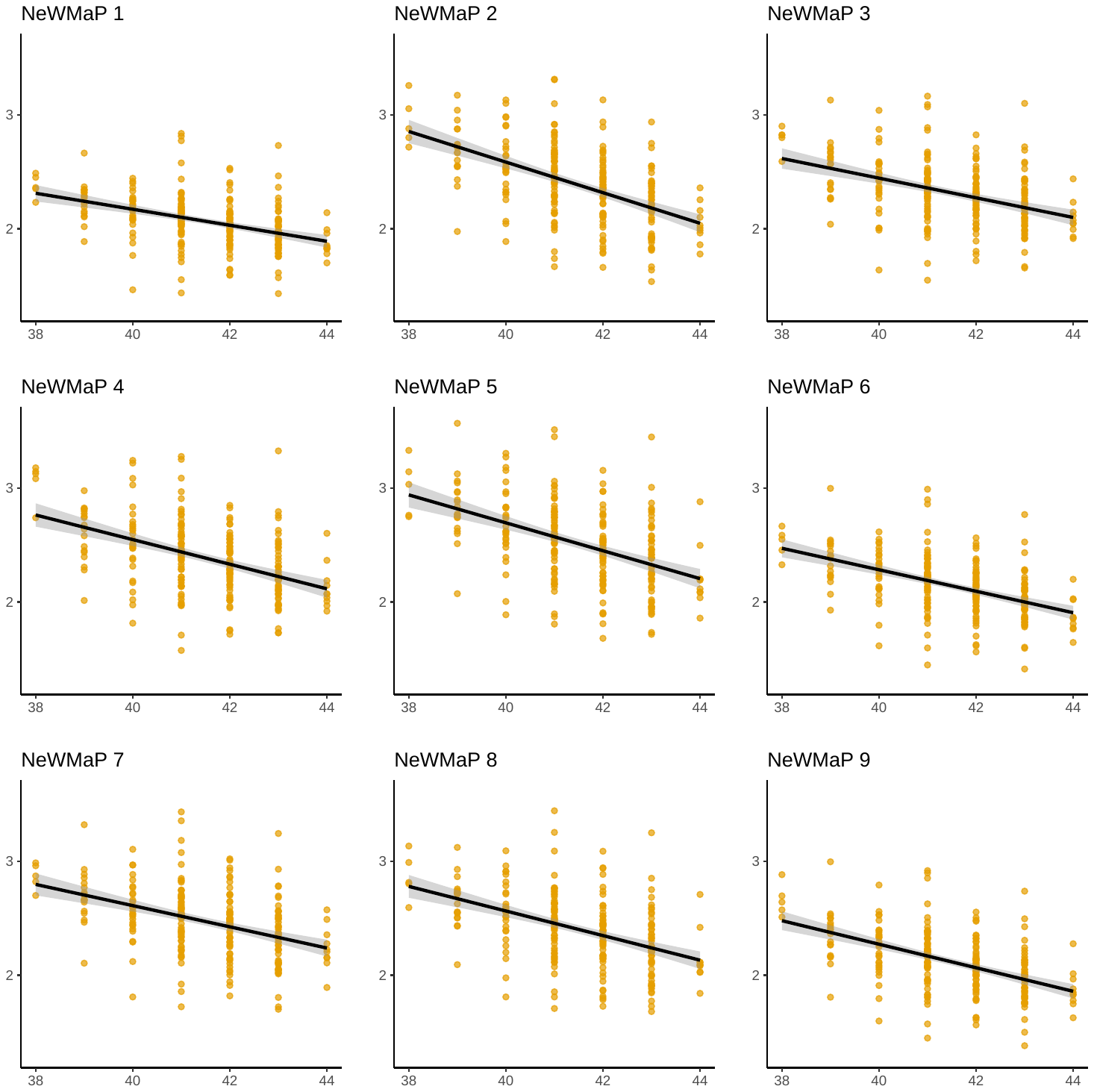

Figure S5. Age-related changes in mean NeWMaP T_2_w/T_1_w signal ratio in the eLABE cohort.

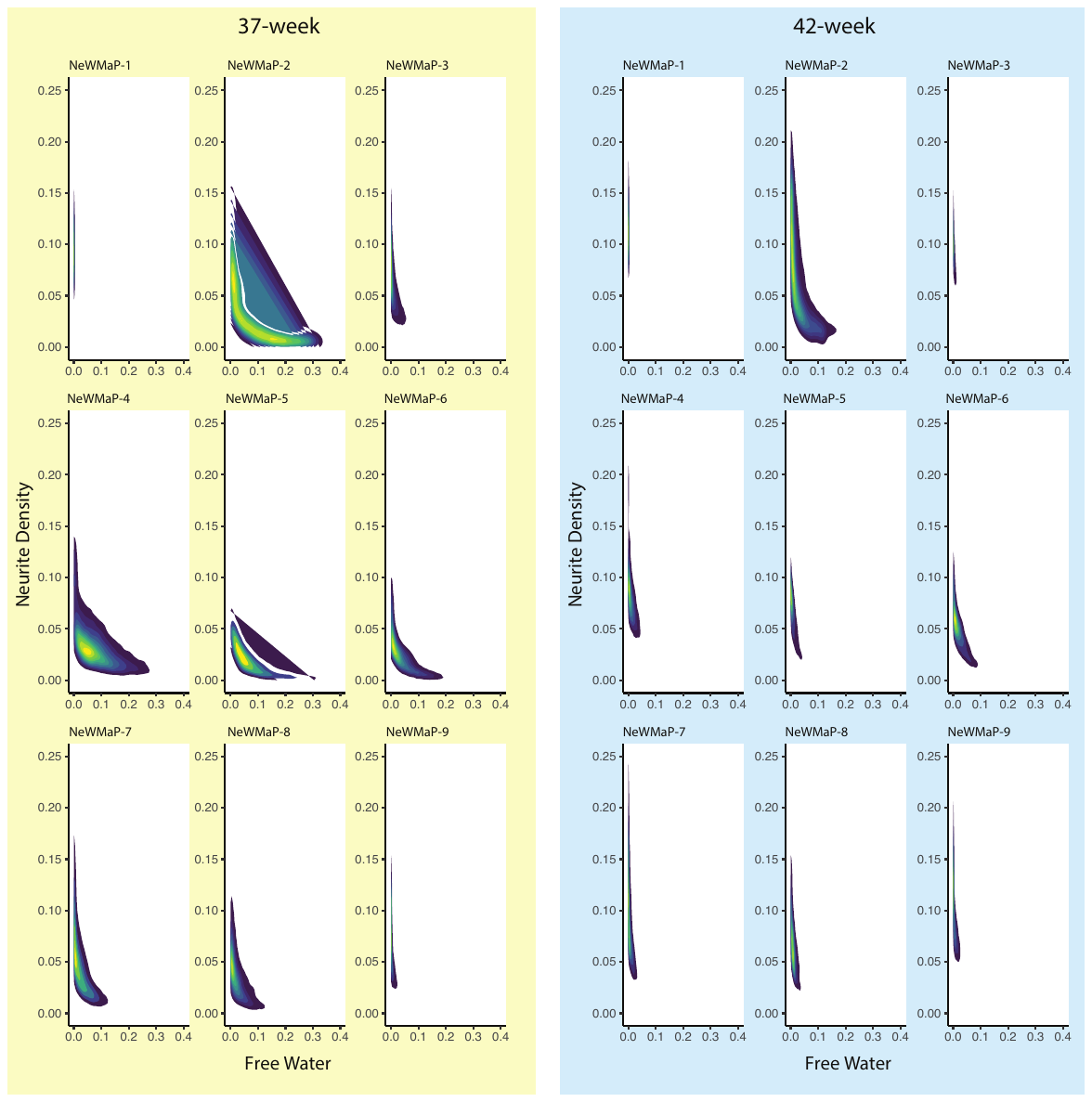

Figure S6. Two-dimensional density plots demonstrating characteristic free water (X-axis) and neurite density (Y-axis) index for each NeWMaP at two age groups (Right: 37-week; Left: 42-week postmenstrual age). These values were extracted from weekly average neurite density and free water maps in the dHCP cohort. While NeWMaP-2, NeWMaP-4, NeWMaP-5, and NeWMaP-6 demonstrate the highest free water contents at 37-week postmenstrual age, NeWMaP-1 and NeWMaP-9 show almost no free water even at this early age. There is a precipitous decline in free water content and moderate increase in neurite density across all NeWMaPs at 42-week postmenstrual age. NeWMaP-2 and to a lesser extent NeWMaP-6 appear to preserve their free water content until 42-week postmenstrual age.

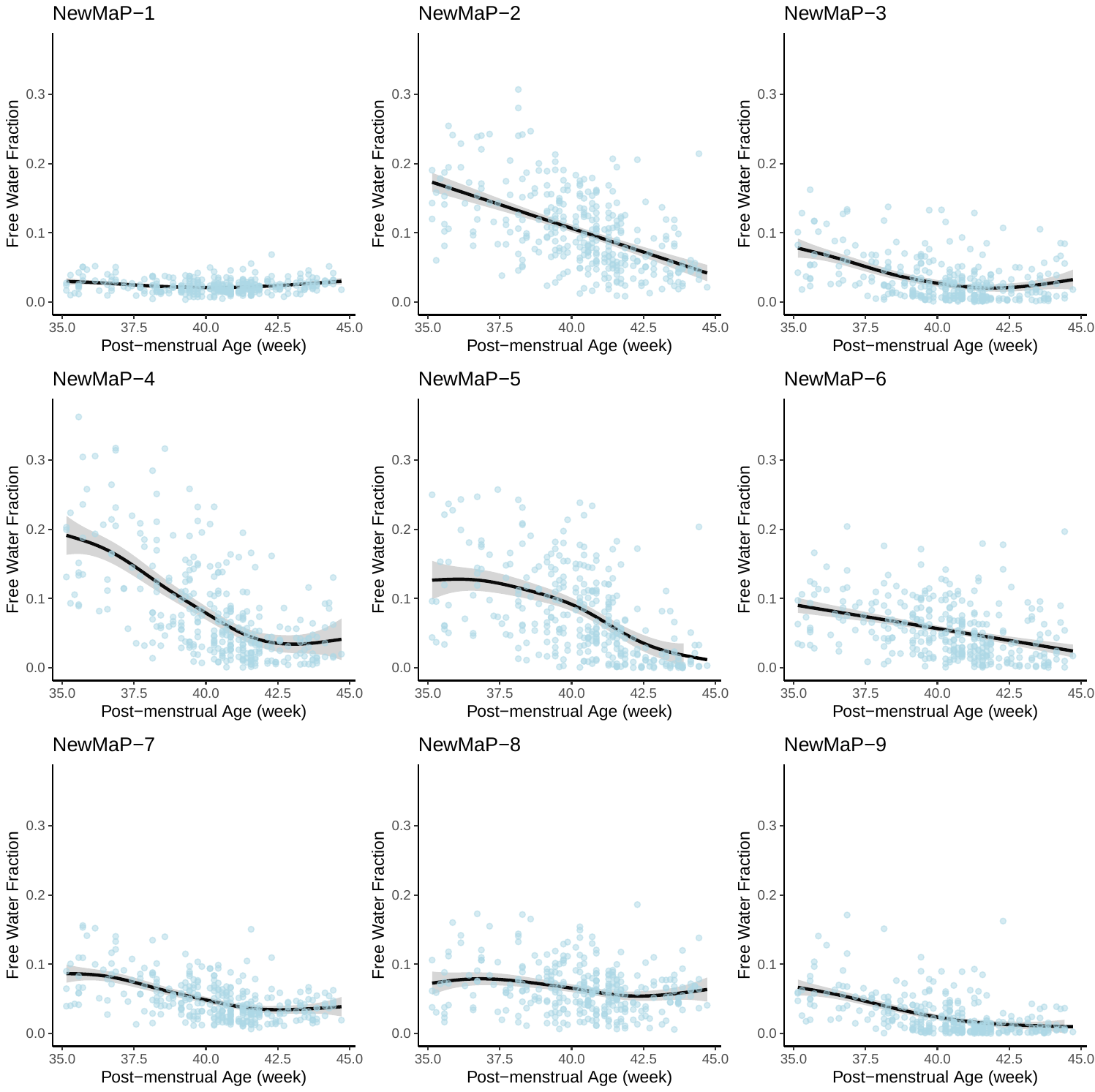

Figure S7. Age-related changes in mean NeWMaP free water content in the dHCP cohort.

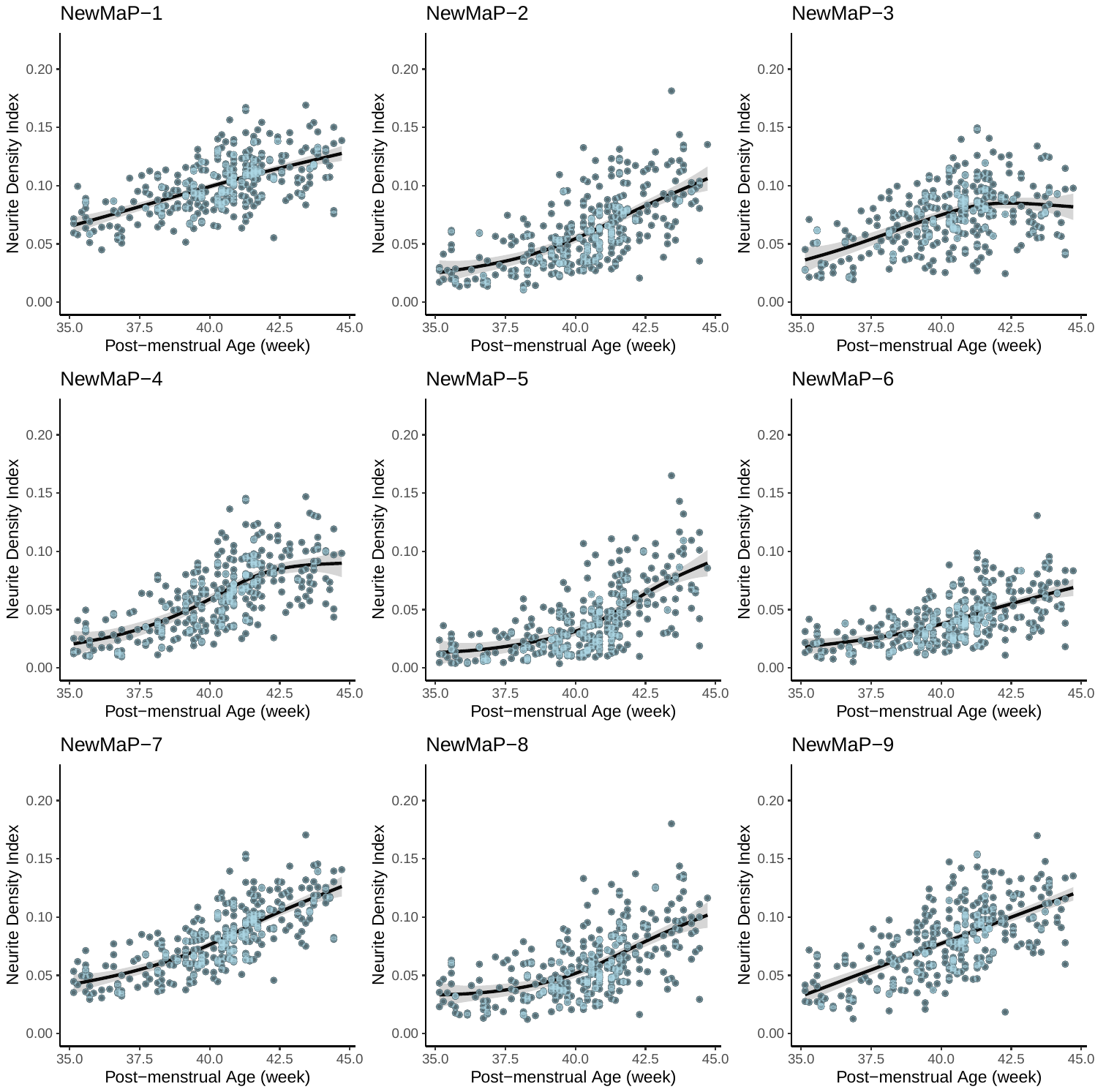

Figure S8. Age-related changes in mean NeWMaP neurite density in the dHCP cohort.

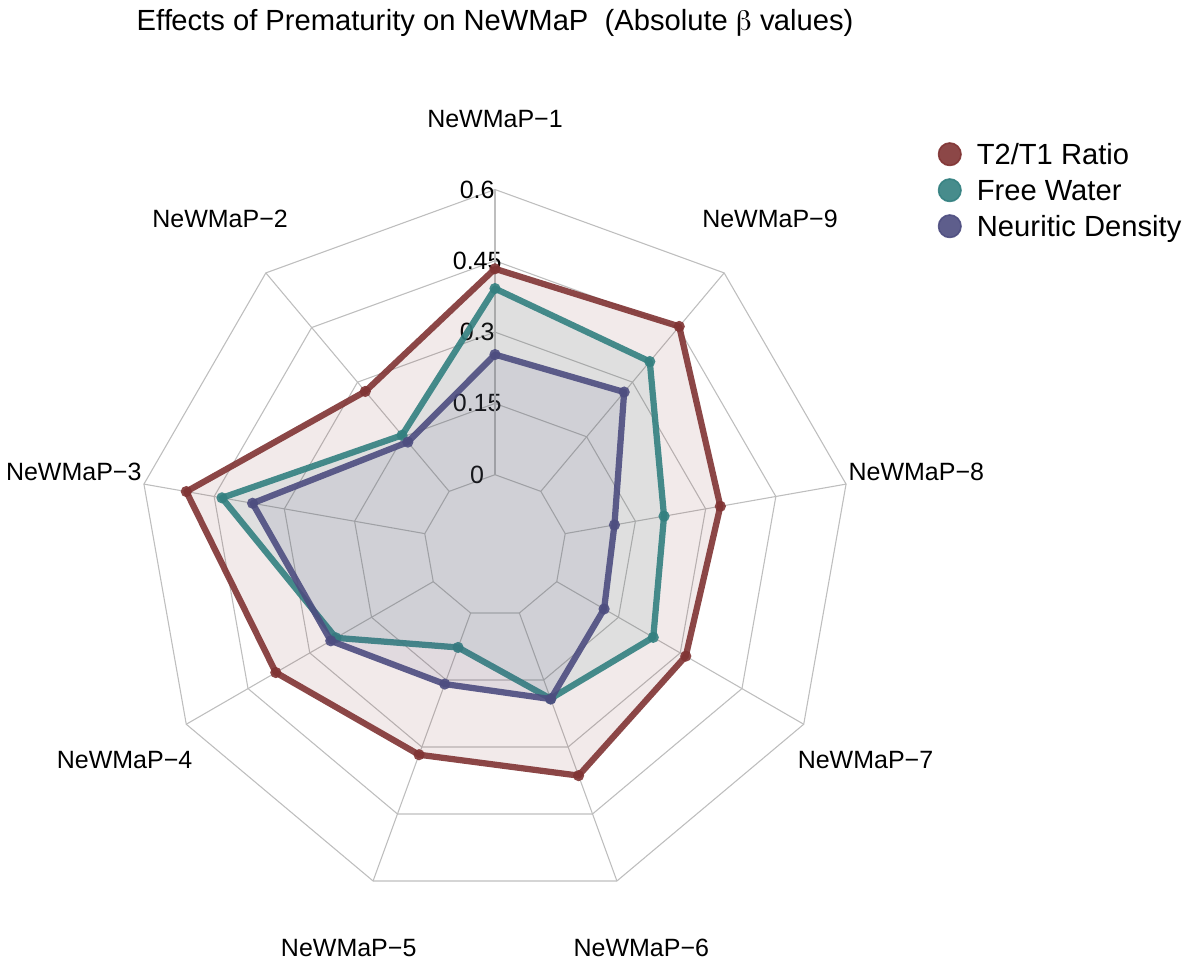

*Figure S9*. Effects of prematurity on mean NeWMaP T_2_w/T_1_w signal ratio, free water content, and neurite density in term-equivalent or older newborns.

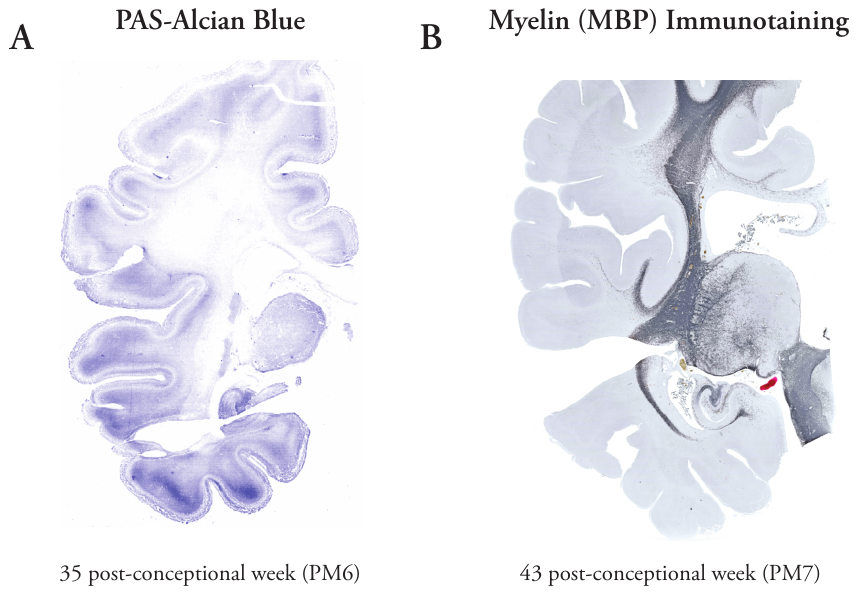

Figure S10. (A) Coronal section through the frontoparietal junction/temporal lobe stained with PAS-Alcian Blue (Alcian Blue stain vector) showing more intense Alcian Blue staining in the gyri (most notable in the temporal lobe). These primarily correspond to superficial white matter NeWMaP-3 and NeWMaP-4. (B) Myelin staining of a coronal section through the frontoparietal junction/temporal immunostained with anti-MBP showing myelination in deep white matter and medial temporal superficial white matter (largely corresponding to NeWMaP-1 and NeWMaP-7).

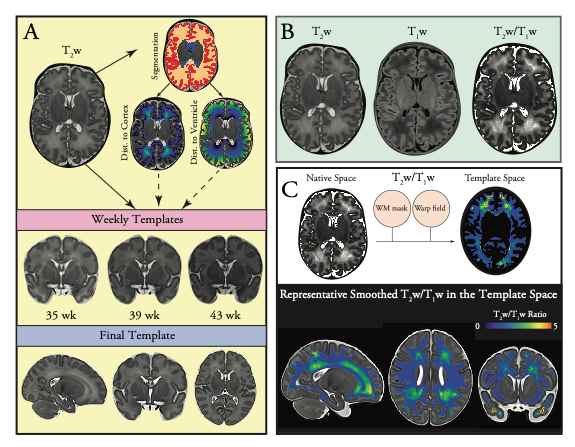

Figure S11. Overview of image preprocessing and registration in the dHCP and eLABE cohorts (performed separately). (A) In the neonatal white matter, distances to the ventricles and to the cerebral cortex convey important topological features that are highly relevant to early postnatal developmental processes. To ensure that these topological features are preserved with nonlinear registrations, we used cortical distance maps (depicting distance to the cerebral cortex in the white matter mask) and ventricular distance maps (depicting distance to the lateral ventricles in the white matter mask) in addition to T2w images for creating templates and image registrations. Given substantial variation in MR signal across different age groups, weekly templates were initially created. The final template was generated by creating a template from the weekly templates (i.e., template of templates). (B) T_2_w/T_1_w ratio maps were registered to the template space using the warp fields. White matter T_2_w/T_1_w ratio maps in the template space were used for further analysis.
